# *Tbx5* maintains atrial identity by regulating an atrial enhancer network

**DOI:** 10.1101/2023.04.21.537535

**Authors:** Mason E. Sweat, Yangpo Cao, Xiaoran Zhang, Ozanna Burnicka-Turek, Carlos Perez-Cervantes, Brynn N. Akerberg, Qing Ma, Hiroko Wakimoto, Joshua M. Gorham, Mi Kyoung Song, Michael A. Trembley, Peizhe Wang, Fujian Lu, Matteo Gianeselli, Maksymilian Prondzynski, Raul H. Bortolin, Jonathan G. Seidman, Christine E. Seidman, Ivan P. Moskowitz, William T. Pu

**Author notes:** Authors contributed equally (Mason E. Sweat, Yangpo Cao).

## Abstract

Understanding how the atrial and ventricular chambers of the heart maintain their distinct identity is a prerequisite for treating chamber-specific diseases. Here, we selectively inactivated the transcription factor *Tbx5* in the atrial working myocardium of the neonatal mouse heart to show that it is required to maintain atrial identity. Atrial *Tbx5* inactivation downregulated highly chamber specific genes such as *Myl7* and *Nppa*, and conversely, increased the expression of ventricular identity genes including *Myl2*. Using combined single nucleus transcriptome and open chromatin profiling, we assessed genomic accessibility changes underlying the altered atrial identity expression program, identifying 1846 genomic loci with greater accessibility in control atrial cardiomyocytes compared to KO aCMs. 69% of the control-enriched ATAC regions were bound by TBX5, demonstrating a role for TBX5 in maintaining atrial genomic accessibility. These regions were associated with genes that had higher expression in control aCMs compared to KO aCMs, suggesting they act as TBX5-dependent enhancers. We tested this hypothesis by analyzing enhancer chromatin looping using HiChIP and found 510 chromatin loops that were sensitive to TBX5 dosage. Of the loops enriched in control aCMs, 73.7% contained anchors in control-enriched ATAC regions. Together, these data demonstrate a genomic role for TBX5 in maintaining the atrial gene expression program by binding to atrial enhancers and preserving tissue-specific chromatin architecture of atrial enhancers.

## Introduction

The cardiac atria rapidly propagate the electrical impulse of the heart beat and coordinately contract to efficiently move blood to the cardiac ventricles^1^. Disrupting these processes results in reduced cardiac pump function and abnormal heart rhythms, including atrial fibrillation (AF), a common and life-shortening arrhythmia^2^. Understanding the pathobiology of atrial diseases such as AF and developing improved treatments requires an in-depth understanding of the regulatory networks that support healthy atrial function.

Atrial and ventricular cardiomyocytes (aCMs and vCMs) are distinct cardiomyocyte subtypes specialized for distinct physiological functions^3–5^. The mechanisms that specify and maintain these differences are only beginning to be elucidated. Factors shown to promote ventricular identity and impede aCM gene expression include FGF, NKX2-5/2-7, IRX4, HEY2, and HRT2^6–11^. In contrast, only a single transcription factor (TF), NR2F2, has been demonstrated to promote atrial identity during early embryonic development^12^. Inactivation of NR2F2 later in heart development did not alter atrial identity^12^, demonstrating its selective role in atrial specification but not maintenance and highlighting that different factors may promote atrial specification versus maintenance.

The cardiac TF *TBX5* is enriched in postnatal aCMs and directly promotes the expression of several atrial specific genes including atrial natriuretic factor (*Nppa)*, connexin 40 (*Cx40*), and bone morphogenic protein 10 (*Bmp10*)^13^*. TBX5* mutations cause congenital cardiac malformations, including atrial septal defect, and upper limb malformations^14^. A familial *TBX5* missense variant, TBX5^G125R^, causes in patients and in overexpressing mice^15, 16^. Widespread postnatal *Tbx5* inactivation in mice using *Rosa26^CreER^* also caused ^17–19^. These data suggest that TBX5 is essential for atrial homeostasis, albeit TBX5’s essential functions in vCMs, conduction system, and nodal tissues potentially confound the interpretation of this widespread inactivation model^20, 21^.

Here, we dissect TBX5’s regulation of atrial gene regulatory networks. Using a model of postnatal, aCM-selective *Tbx5* inactivation, we demonstrate alteration of aCM structure and rapid development of .Concurrent single nucleus transcriptome and open chromatin sequencing (snRNAseq and snATACseq) coupled with H3K27ac HiChIP showed that aCMs lacking *Tbx5* lose accessibility and chromatin looping at many enhancers of aCM-selective genes.

## RESULTS

### Nppa-Cre specifically inactivates Tbx5 in aCMs

To inactivate a loxP-flanked *Tbx5* allele (*Tbx5^flox^*) selectively in postnatal aCMs, we expressed Cre using the *Nppa* promoter and the cardiotropic adeno-associated virus serotype 9 (AAV9), which was previously shown to selectively recombine floxed alleles in atria^22, 23^. Since its activity in sino-atrial or atrioventricular nodes (SAN and AVN) had not previously been assessed, we administered AAV9:*Nppa-EGFP* at postnatal day P8 (P8) to *Hcn4^CreER/+^*; *Rosa26^LSL:Tomato^* mice. At P21 and P22, tamoxifen was administered to activate tomato expression in the conduction system including the sinoatrial and atrioventricular nodes (**Supp. Fig. 1a**). Hearts exhibited strong GFP signal in the atria and non-overlapping Tomato signal positioned consistent with the SAN (**Supp. Fig. 1b,b’**). Cryosections revealed mutually exclusive Tomato fluorescence in SAN and AVN and GFP in adjacent working aCMs (**Supp. Fig. 1c, boxes 1+2**). These data demonstrate that AAV9 and the *Nppa* promoter express in aCMs and not SAN or AVN..

Next we characterized AAV9:Nppa-Cre inactivation of *Tbx5*. Crossing *Tbx5^Flox^* mice to the Cre-inducible H2b-mCherry reporter line (*Rosa^H2B-mch^*) allowed for the identification of myocytes transduced by AAV9:Nppa-Cre. At P2, *Tbx5^+/+^, Rosa^H2B-mch^; Tbx5^Flox/+^, Rosa^H2B-mch^;* and *Tbx5^Flox/Flox^, Rosa^H2B-mch^* mice were treated with AAV9:*Nppa-Cre*. At P20, H2B-mCherry fluorescence was localized to the atria, and AAV9:Nppa-Cre *Tbx5^Flox/Flox^* atria were markedly enlarged (**Fig. 1b-d**). In histological sections, aCMs but not vCMs exhibited prominent nuclear mCherry signal (**Fig. 1e-g**). Immunoblots revealed loss of TBX5 protein in atrial lysates from *Tbx5^Flox/Flox^* mice compared to control Tbx5^+/+^ mice, and no reduction in ventricular lysates (**Fig. 1h**). TBX5 was partially reduced in Tbx5^Flox/+^ mice, consistent with heterozygosity. RT-qPCR confirmed 80% reduction of Tbx5 transcripts in *Tbx5^Fox/flox^* atria and no significant change in *Tbx5^Fox/flox^* ventricles (**Fig. 1i**). Taken together, these results demonstrate AAV9:*Nppa-Cre* effectively and selectively inactivates *Tbx5* in aCMs. Hereafter we refer to *AAV9:Nppa-Cre* treated *Tbx5^flox/flox^* as *Tbx5^AKO^* (aCM knockout).

**Figure 1.**
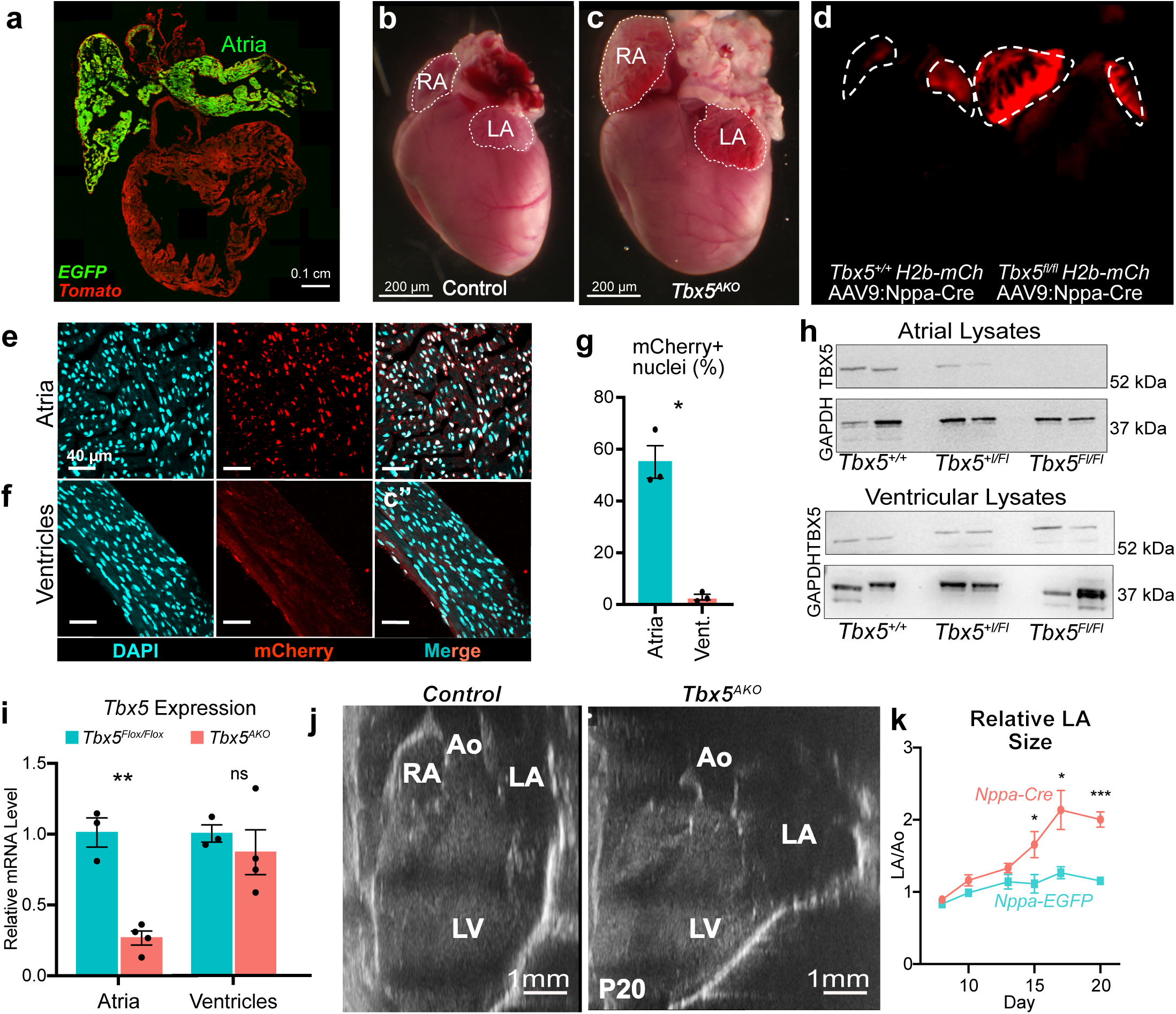
Inactivation of *Tbx5* in aCMs results in atrial remodeling. **a,** Rosa^mTmG^ Cre reporter mice were injected with 2 x10^11^ VG/g AAV9:Nppa-Cre at P2. Hearts were analyzed at P20. Cre-activated EGFP signal was restricted to the atria. **b-d,** Whole mount images of control and atrial-specific *Tbx5* knockout hearts. *Tbx5^+/+^; H2b-mCh* or *Tbx5^Flox/Flox^; H2b-mCh* mice were treated with AAV9:Nppa-Cre at P2. *Tbx5^flox/flox^* mice injected with AAV9:Nppa-Cre (*Tbx5^AKO^)* exhibited enlarged atria. b-c, brightfield. d, red fluorescence channel. **e-g,** AAV9:Nppa-Cre mediated recombination in atria and ventricles. H2B-mCh Cre reporter mice were treated with AAV9:Nppa-Cre at P2. P20 sections of atria (d) and ventricles (e) were analyzed by confocal imaging. Quantification of the percentage of mCherry+ nuclei in atria and ventricles (f). Unpaired *t*-test, *, P = 0.0123. n=3 hearts per group. **h,** TBX5 protein level in *Tbx5^AKO^* and control atria and ventricles. *Tbx5^+/+^, Tbx5^+/flox^*, and *Tbx5^Flox/flox^* mice were treated with AAV9:Nppa-Cre at P2. Atrial and ventricular lysates, prepared at P20, were analyzed by western blotting to detect TBX5 and GAPDH. **i,** RT-qPCR for *Tbx5* mRNA from atria and ventricles. Unpaired t-test: **, P = 0.0079; ns, not significant (P=0.48). n=3-4 hearts per group. **j-k,** Echocardiographic assessment of atrial size. i, representative echo images used to quantify atrial size, compared to aortic size. RA, right atrium; LA, left atrium; LV, left ventricle; Ao, aorta. j, Relative LA size of control and *Tbx5^AKO^*left atria, normalized to aortic diameter. Unpaired t-test: *, P<0.05; ***, P=0.0001. n=4 *Tbx5^AKO^* and 5 control mice. Error bars represent SEM.

### Atrial remodeling and fibrillation in Tbx5^AKO^ mice

To further examine the atrial phenotype of *Tbx5^AKO^ mice,* we performed serial echocardiography from P8 to P20 (**Fig. 1j**). After P15, the atria were significantly dilated in *Tbx5^AKO^* compared with controls treated with *AAV-Nppa-EGFP*, as assessed by the ratio of left atrium to aorta diameters (**Fig. 1k**). There was no significant difference in ventricular function measured at P20 between control and *Tbx5^AKO^* mice (**Supp. Fig. 2f**). We assessed fibrosis by collagen staining (Masson’s trichrome) and RT-qPCR for fibrosis genes *Col1a1* and *Postn* (**Supp. Fig. 2a, b**). In contrast to some other models of that demonstrated fibrotic remodeling of the atrial chambers^22, 23^, inactivating *Tbx5* did not result in fibrosis at this early time point.

We assessed aCM morphology by immunostaining for the Z-line protein sarcomeric α-actinin (SAA) and the junctional sarcoplasmic reticulum protein FSD2^24^. In control aCMs and aCMs with one functional copy of *Tbx5*, we observed the expected striated Z-line staining pattern (**Supp. Fig. 2c-c’,d-d’**). However, in aCMs lacking both copies of *Tbx5,* this striated pattern was absent in many myocytes, as illustrated by plotting SAA signal intensity along the aCM long axis (**Supp. Fig. 2e**).

Since widespread *Tbx5* inactivation caused AF^17–19^, we obtained electrocardiograms (EKGs) on *Tbx5^AKO^*mice at P21. In *Tbx5^Flox/Flox^* mice treated with *AAV9:Nppa-GFP* (control), the EKG showed P waves preceding each QRS complex and similar intervals between QRS complexes (RR intervals), consistent with normal sinus rhythm. This regularity of RR intervals was illustrated by the tight clustering of points in “Poincaré plots” of the RR interval of two successive beats (**Supp. Fig. 2h**). In *Tbx5^AKO^* animals, there was a drastic change in EKG morphology, including the loss of P waves and irregularly irregular RR intervals (**Supp. Fig. 2g**), hallmarks of AF. This irregularity was reflected by dispersion of points in the Poincaré plots of Tbx5^AKO^ mice. We quantified the irregularity by calculating the standard deviation of the RR interval (SDRR) at P21. Strikingly, the mean SDRR was 5 times greater in *Tbx5^AKO^* animals compared to controls (**Supp. Fig. 2i**). Serial measurements between P8 and P21 showed significant elevation of SDRR between P14 and P17 (**Supp. Fig. 2j**), indicating that *Tbx5* is required in aCMs to maintain atrial rhythm even in young mice.

### Tbx5 promotes aCM identity

Recent work aimed at identifying important regulators of chamber identity uncovered the TBOX motif as an enriched motif in atrial-specific enhancers and demonstrated elevated expression of *Tbx5* in aCMs compared to vCMs^3^. Other studies demonstrated that TBX5 is required for expression of atrial-specific genes, including *Nppa* and *Bmp10*^25^. To further evaluate the requirement of *Tbx5* for atrial identity, we stained for MYL7, the atrial specific myosin regulatory light chain, in P21 heart sections from untreated *Tbx5^Flox/Flox^; Rosa^H2B-mch^ and* AAV9:Nppa-Cre-treated *Tbx5^Flox/Flox^; Rosa^H2B-mch^*, or *Tbx5^Flox/+^* mice. MYL7 immunoreactivity similarly localized in Cre-treated *Tbx5^Flox/+^*and untreated *Tbx5^Flox/Flox^* aCM A-bands, demonstrating that *Tbx5* heterozygosity is sufficient to support aCM expression of MYL7 (**Fig. 2a**). In contrast, Cre-treated *Tbx5^AKO^* aCMs lacking *Tbx5* (mCherry+) lost MYL7 staining almost entirely (**Fig. 2a, arrows**). Within the same tissue, neighboring aCMs that did not undergo Cre recombination (mCherry-) retained MYL7, indicating that this phenotype is cell autonomous and not secondary to tissue level abnormalities such as AF.

**Figure 2.**
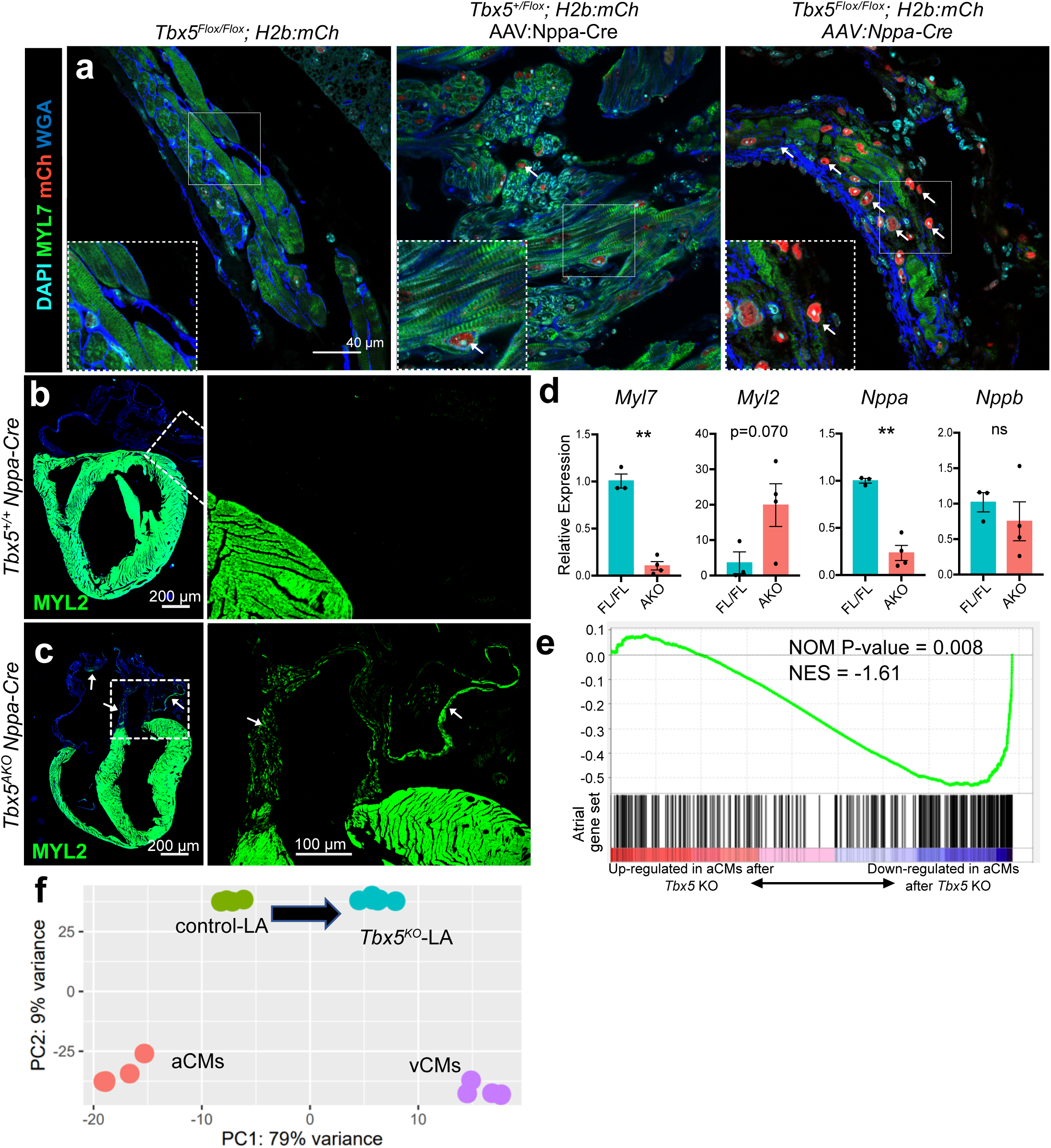
Inactivating *Tbx5* alters the expression of atrial identity genes. **a,** Staining for MYL7, mCh and WGA in the left atrium of uninjected *H2b:mCh Tbx5^Flox/Flox^;* AAV9:Nppa-Cre *H2b:mCh, Tbx5^Flox/+^;* and AAV9:Nppa-Cre, *H2b:mCh Tbx5^AKO^* mice. Arrows in the middle and right panels denote mCherry+ cells, which were transduced by AAV9:Nppa-Cre. White boxed regions magnified in insets in the lower left corners. **b-c,** MYL2 expression staining in *Tbx5^AKO^* atria. *Tbx5^+/+^* and *Tbx5^Flox/Flox^* mice were treated with AAV9:Nppa-Cre. Paraffin sections were stained for ventricular marker MYL2. White boxed regions are enlarged to right. Arrows in **c** indicate *Tbx5^AKO^* atrial regions that express MYL2 in atria, which were not observed in control atria. **d,** Measurement of aCM and vCM selective transcripts in control and *Tbx5^AKO^* atria. *Myl7, Myl2, Nppa,* and *Nppb* transcripts were measured from pooled atrial samples of the indicated genotype. Error bars indicated SEM. Unpaired t-test: *, P<0.05; **, P<0.01. ns, not significant. n=3-4 per group. **e,** Enrichment of aCM-selective genes among genes downregulated in *Tbx5^AKO^* aCMs. Gene set enrichment analysis was used to test genome-wide effect of Tbx5 atrial knockout on expression of an atrial gene set containing genes significantly upregulated in aCMs compared to vCMs. **f,** Genome-wide effect of *Tbx5* atrial knockout on expression of left atrial genes. Principle component analysis was used to compare transcriptomes of isolated aCMs, isolated vCMs, control LA, and Tbx5 knockout LA (from GSE 129503). The black arrow highlights the shift of aCM gene expression towards vCMs along PC1 after inactivating *Tbx5*.

The myosin regulatory light chain expressed in vCMs is MYL2, rather than aCM-specific MYL7. In AAV9:Nppa-Cre-treated *Tbx5^+/+^* heart sections, MYL2 immunoreactivity was robust in vCMs and not detected in aCMs (**Fig. 2b**). In contrast, MYL2 expression was detected in *Tbx5^AKO^* aCMs (**Fig. 2c**). RT-qPCR confirmed downregulation of aCM-specific transcripts *Myl7* and *Nppa* and upregulation of vCM-specific transcript *Myl2* in *Tbx5^AKO^* aCMs. However, there was no significant change in the aCM-specific transcript *Nppb* (**Fig. 2d**).

To better understand changes in the atrial gene expression profile, we examined bulk RNA-seq of atrial tissue from a previous study of adult ubiquitous *Tbx5* knockout^17^. We compared differentially expressed left atrial genes to sets of genes with aCM-selective or vCM-selective expression as defined by RNA sequencing of purified cardiomyocytes from each chamber (|fold change| > 1.5, Padj < 0.05)^3^. Gene set enrichment analysis with aCM-selective genes revealed statistically significant reduction in *Tbx5* knockout atria (**Fig. 2e**). This effect was also seen in principal component analysis (PCA) plots of aCM, vCM, control left atria, and adult ubiquitous Tbx5 knockout left atria, in which *Tbx5*-ablated atrial gene expression shifted away from aCMs and towards vCMs on principle component 1 (**Fig. 2f**). Together, these data demonstrate that *Tbx5* promotes the expression of a subset of atrial specific genes and represses the expression of a subset of ventricular genes.

### Tbx5 overexpression promotes atrial identity in ventricular cardiomyocytes

To determine if *Tbx5* is sufficient to promote atrial identity, we turned to a model of *Tbx5* overexpression in which cardiomyocyte specific Myh6-Cre activated a Cre-dependent *Tbx5* transgene^26^ (*Tbx5-OE*; **Supp. Fig. 3a**). Bulk RNA-sequencing of *Tbx5-OE* and control left ventricular myocardium at 6 weeks of age revealed thousands of significantly altered transcripts (**Supp. Fig. 3b,c; Supp. Table 1-2**). TBX5 overexpression significantly upregulated 178 aCM-selective genes in ventricular myocardium, compared to only 69 that were downregulated (**Supp. Fig. 3d**; Fisher exact test: P< 0.00001). Furthermore, TBX5 overexpression downregulated 181 vCM-selective genes, compared to only 73 that were upregulated (**Suppl. Fig. 3e**; Fisher exact test: P<0.00001). These data demonstrate that *Tbx5* overexpression is sufficient to drive the transcriptome of established vCMs towards an aCM profile, activating 15.8% of all aCM-selective genes while repressing 20.7% of vCM-selective genes.

### Concurrent snRNA- and snATAC-seq analysis of Tbx5^AKO^ and control left atrial tissue

To more comprehensively evaluate the effects of Tbx5^AKO^ on cell composition and gene expression, left atria were isolated from P21 *Tbx5^flox/flox^* animals treated with *AAV9:Nppa-GFP* (control) or *AAV9:Nppa-Cre* and used as input for single nucleus multiome (RNA and ATAC sequencing) analysis (**Fig. 3a**). In total, 14,583 nuclei passed rigorous quality and doublet filters, and an average of 1,926 genes were detected per nucleus (**Supp. Fig. 4c, Supp. Table 1**). ATAC fragments from each of the samples demonstrated the expected enrichment at the transcription start site (TSS) of mouse genes (**Supp. Fig. 4b**). UMAP embeddings were created separately for the transcriptome and accessibility assays (**Fig. 3b,c**), and using a weighted nearest neighbors (WNN) approach to integrate both datasets (**Fig. 3d**)^27–29^. Biological duplicate *Tbx5^AKO^*and control samples showed excellent overall overlap of each replicate (**Supp. Fig. 4**), demonstrating reproducibility of the data. Clustering identified 20 cell states. The identities for these clusters were inferred using marker genes specific for different cardiac cell types. Six clusters expressed cardiomyocyte markers *Tnnt2, Ryr2*, and *Mybpc3* (**Fig. 3e**). Remarkably, control and *Tbx5^AKO^* aCMs largely clustered separately, with myocyte clusters 1-4 predominantly comprising control aCMs and clusters 5-6 predominantly containing *Tbx5^AKO^* aCMs.

**Figure 3.**
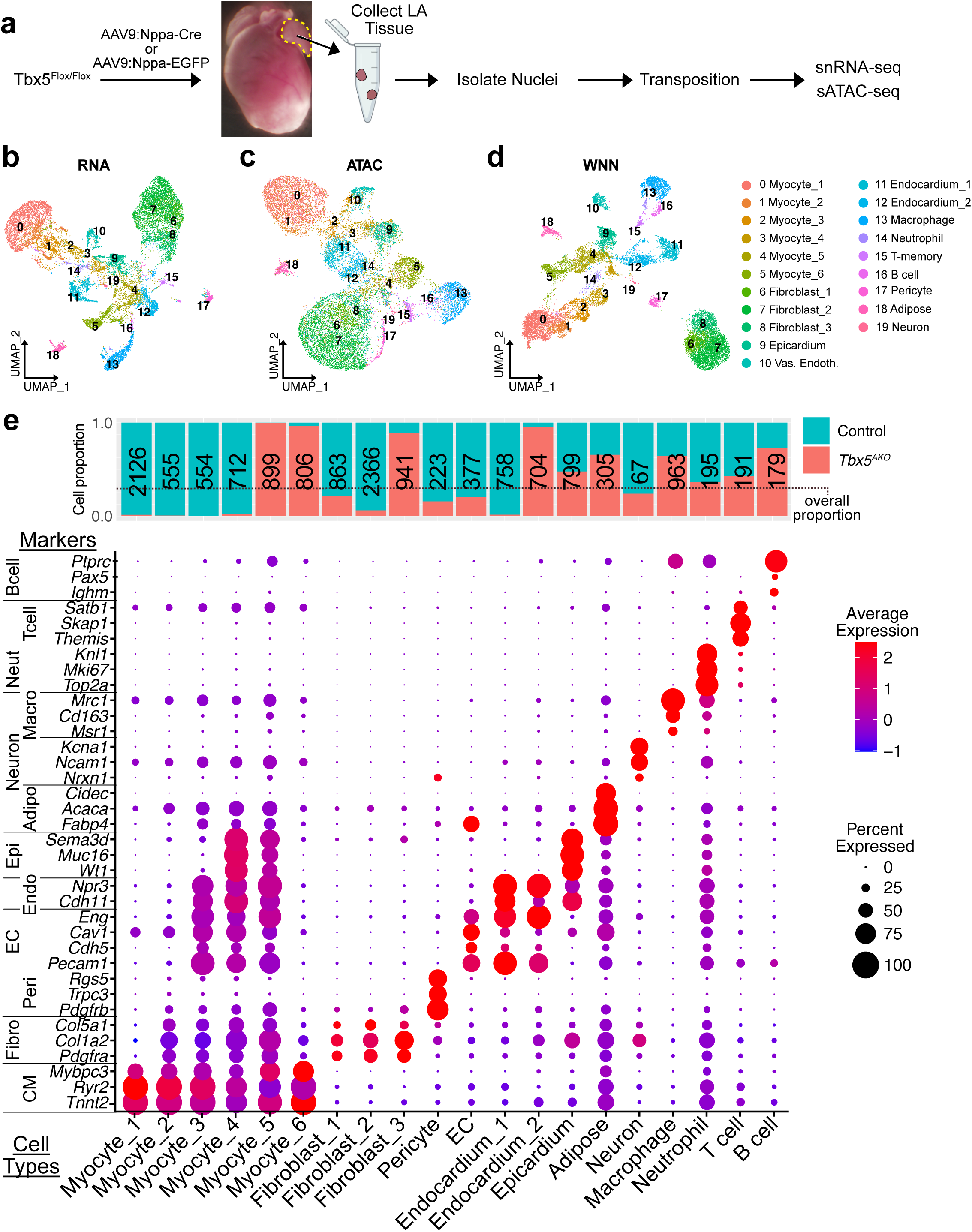
Concurrent scRNA-seq and scATAC-seq analysis of cell states in control and *Tbx5^AKO^*atria. **a,** Experimental design. Neonatal Tbx5^Flox/Flox^ mice were treated with AAV9:Nppa-EGFP (control) or AAV9:Nppa-Cre (KO). At P20, left atria were dissected from two mice per replicate for the analysis. Isolated nuclei were used for concurrent scATAC-seq and scRNA-seq. Two replicates were prepared per group. **b-d,** Clustering of cell states based on scRNA-seq, scATAC-seq, or both assays using weighted nearest neighbor (WNN) analysis. **e,** Cluster identities were established using cell-type specific markers. The total number of nuclei as well as the contribution from control and AKO samples are shown. The black dotted line indicates the overall relative contribution of KO and control nuclei to the assay. Other dataset metrics are available in **Supplemental Figure 2**. CM, cardiomyocyte; Fibro, fibroblast; Peri, pericyte; EC, endothelial cell; Endo, endocardium; Epi, epicardium; Adipo, adipocyte; Macro, macrophage; Neut, neutrophil.

To understand the relationship between the myocyte clusters, we performed trajectory analysis^30^. Control clusters (Myocyte_1 to Myocyte_4) and *Tbx5^AKO^* clusters (Myocyte_5 and Myocyte_6) arranged into separate trajectories (**Fig. 4a; Supp. Fig. 5a, b**). Within each trajectory, comparison of the cluster at the end versus the start (Myocyte_1 vs. Myocyte _4 for control and Myocyte_6 vs. Myocyte_5 for *Tbx5^AKO^*) revealed similar gene expression changes, including increased expression of genes associated with sarcomere integrity (*Ttn, Myh6*) and contraction (*Ryr2, Atp2a2)* (**Supp. Fig. 5c, d**). GO term analysis of these differentially expressed genes demonstrated similar enrichment in the late pseudotime clusters of terms related to muscle contraction and sarcomere organization. Terms enriched in the early pseudotime clusters were not specific for cardiomyocytes and included extracellular matrix organization, regulation of cell migration, and vascular development. Taken together with the trajectory analysis (**Fig. 4a**), these results indicate that the multiome analysis captured aCMs at different stages of maturity, with Myocyte_1 and Myocyte_6 representing the most mature clusters of control and *Tbx5^AKO^*aCMs, respectively. Unless stated otherwise, future analyses are primarily focused on the mature Myocyte_1 and Myocyte_6 clusters, which we refer to as the “control aCM” and “KO aCM” clusters, respectively.

**Figure 4.**
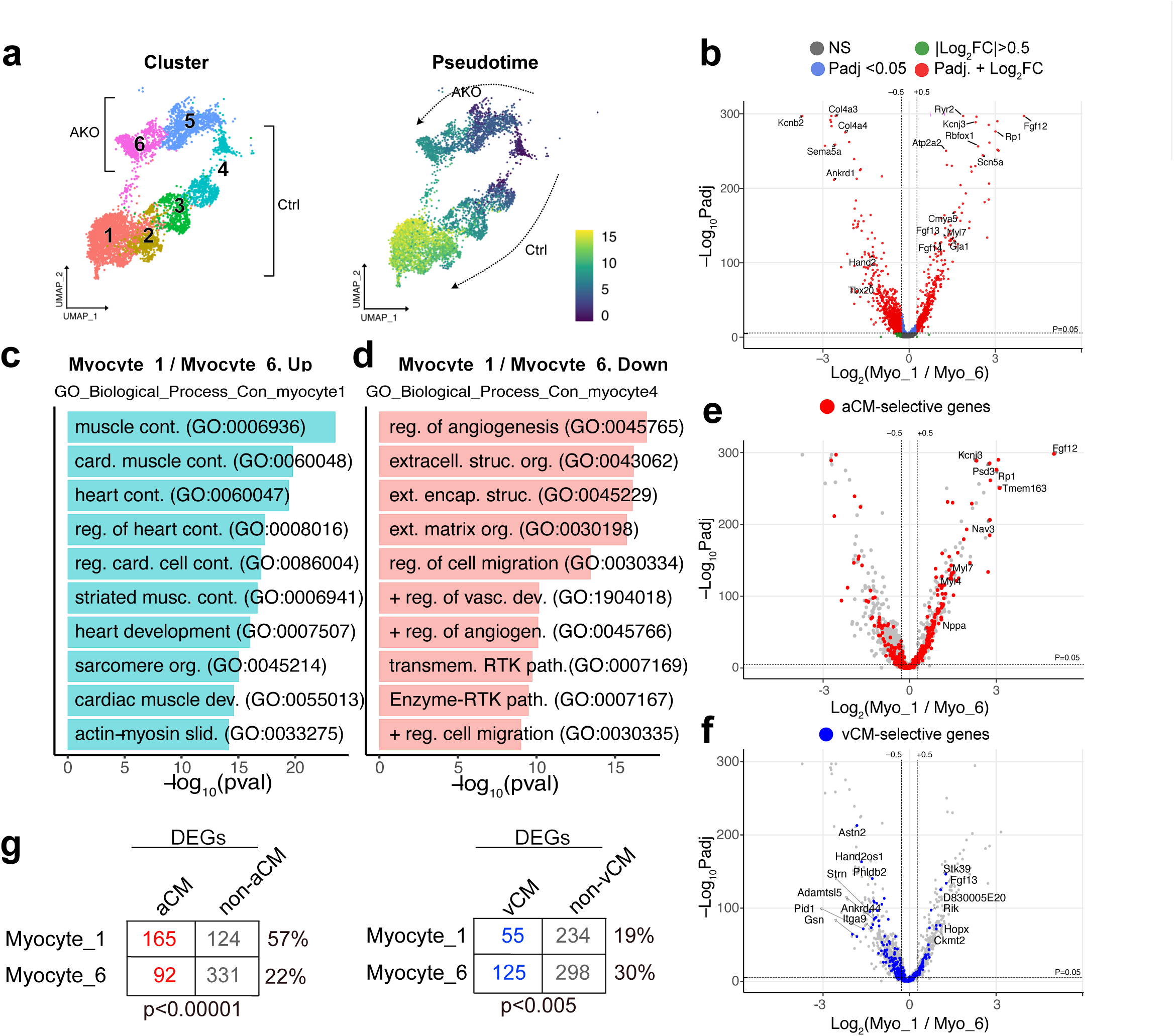
TBX5 is required to promote the expression of aCM genes. **a,** Pseudotime trajectory of myocyte clusters. Trajectory analysis was performed with Monocle3. Two separate trajectories were inferred. Clusters containing predominantly control (Ctrl) and knockout (AKO) myocytes participated in separate trajectories. Clusters 1 (control) and 6 (KO) were the endpoints of each trajectory. **b,** Differential gene expression analysis was performed, comparing the clusters located at the end of the control trajectories (Myocyte_1, control and Myocyte_6, KO). The top most differentially expressed genes (DEGs) are labeled. **c-d,** Biological process GO terms enriched for genes with significantly greater expression in control (blue, Myocyte_1)) or KO (pink, Myocyte_6) clusters. The top 10 terms terms are shown. **e-g,** Distribution of chamber-selective genes among genes differentially expressed between control or KO myocyte clusters (Myo_1 vs Myo_6). aCM-selective genes (e) and vCM-selective genes (f) were disproportionately found in genes with significant upregulation in control (Myo_1) or KO (Myo_6), respectively. Statistical analysis of distribution of aCM genes (left) or vCM genes (right) among Myo_1 vs Myo_6 DEGs. Significance of 2 x 2 tables were evaluated using Fisher’s exact test.

Next we identified gene expression differences between the control aCM and KO aCM clusters (**Supp. Table 3**). Differentially expressed genes (DEGs) more highly expressed in the control aCM cluster included ion channel genes *Scn5a*, *Atp2a2*, and *Ryr2*, known direct TBX5 targets which are critical for action potential propagation and calcium handling^31, 32^ (**Fig. 4b**). *Sbk2*, a gene recently shown to be essential for atrial sarcomere integrity^33^, was also more highly expressed in control compared to *Tbx5^AKO^* aCMs, possibly contributing to the observed disorganization of *Tbx5^AKO^* sarcomeres (**Supp. Fig. 2C-D**). In the KO aCM cluster, there was an upregulation of structural genes and genes involved with stress, including *Col4a3*, *Col4a4*, and *Ankrd1*. *Ank3*, previously linked with Brugada syndrome, was also upregulated in *Tbx5^AKO^*aCMs^34^. Genes more highly expressed in the control aCM cluster were associated with heart contraction and cardiac conduction, including regulation of the atrial action potential, a known *Tbx5-*dependent feature^17^ (**Fig. 4c,d**).

To further investigate the role of *Tbx5* in atrial identity, we analyzed the distribution of aCM-selective and vCM-selective genes among genes differentially expressed in the control and KO aCM clusters (**Fig. 4e,f**). Strikingly, 57% of the genes enriched in the control aCM cluster were aCM selective, compared to only 22% of genes in the KO aCM cluster (**Fig. 4e,g**; Fisher’s exact test: P<0.00001). Conversely, vCM-selective genes were enriched in the KO aCM clusters (30%, versus 19% of genes more highly expressed in control aCMs; **Fig. 4f,g**; Fisher’s exact test: P=0.005). These data demonstrate that *Tbx5* is necessary to maintain aCM identity.

We compared the DEGs found in *Tbx5^AKO^* in aCMs to DEGs found in TBX5-OE vCMs to determine if TBX5 regulated a common set of chamber-selective genes in the atrium and ventricle (**Supp. Fig. 3f,g**). Overlapping these data showed that TBX5 promoted the expression of a common subset of 45 aCM genes in both chambers, representing 27.3% of the overall aCM genes enriched in the control aCM cluster and 27.1% of the aCM genes enriched in *Tbx5-OE* LV. Of the twenty genes with the greatest shared enrichment in the control aCM cluster and *Tbx5-OE* LV, twelve were aCM genes. In total, TBX5 regulated a total of 286 (over 25%) of all aCM genes, demonstrating the extensive dependence of atrial identity on TBX5.

### TBX5 maintains accessibility at TBX5 binding sites in atrial cis-regulatory elements

Cellular identity is regulated by the interaction of TFs with distal regulatory elements that control gene expression^35^, and we hypothesized that TBX5 exerts its control over atrial identity by binding and modulating the activity of atrial enhancers. To test this hypothesis, we compared chromatin accessibility between control and *Tbx5^AKO^* aCM clusters using the multiome’s scATAC-seq data (**Fig 5a**). For pairwise comparisons within or between start and end points of the control and *Tbx5^AKO^* trajectories, we quantitatively compared accessibility signals between clusters to define differentially accessible regions (Padj < 0.05; **Fig. 5a-b; Supp. Table 4**). Early and late control myocyte clusters had the most differentially accessible regions (5818 regions). In the *Tbx5^AKO^* trajectory, less than 20% (1193) of this number of regions were differentially accessible. These results suggest that maturational changes in both control and knockout involve substantial changes in chromatin accessibility and cis-regulatory element usage, with many dependent on *Tbx5*.

**Figure 5.**
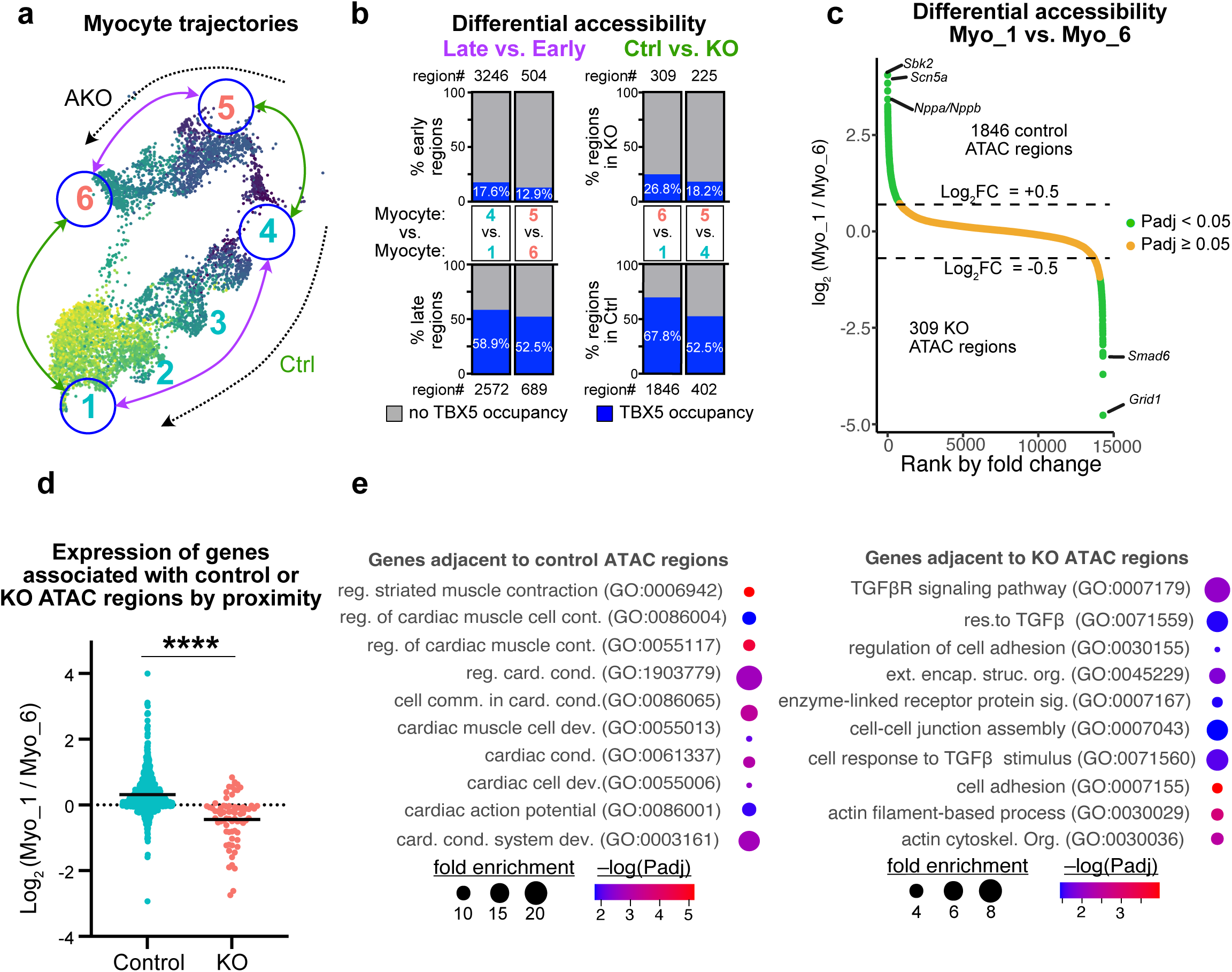
Multiomics reveals a set of *Tbx5-*dependent CREs that promote aCM gene expression. **a,** Myocyte trajectories and comparisons performed to identify differentially accessible genomic regions. Arrows indicate the four pairwise comparisons between the clusters at the start and end of the control and *Tbx5^AKO^*trajectories. **b,** Differentially accessible regions were analyzed for four pairwise comparisons. The number of differentially accessible regions (region #) is shown at the end of each bar. The bar indicates the fraction of these regions that are occupied by TBX5 in wild-type aCMs. **c,** Accessibility change between the control and TBX5 KO aCM clusters. Plotting the average log_2_FC accessibility of regions in the control aCM cluster compared to the KO aCM cluster revealed 1846 significant control regions and 309 significant KO regions (Padj. < 0.05). Genes nearest to the most enriched control and KO regions are listed. **d**, Expression of genes associated with control or KO regions. Control or KO regions were associated with the nearest gene. The expression ratio of each set of genes between Myocyte_1 and Myocyte_6 was compared using the Mann-Whitney test. ****, P<.0001. **e,** GO terms for genes nearest to control regions or KO regions. Genes next to control regions tended to be associated with cardiomyocyte function, while genes proximal to KO regions were associated with TGFβ signaling and remodeling terms.

Comparison between control and *Tbx5^AKO^* clusters at early (Myocyte_4 vs. Myocyte_5) or late (Myocyte_1, the control aCM cluster, vs. Myocyte_6, the KO aCM cluster) pseudotimes identified chromatin accessibility that was dependent upon *Tbx5* (**Fig. 5b**). We integrated these data with TBX5 genomic occupancy in wild-type aCMs, based on high affinity streptavidin pulldown of an endogenously biotinylated *Tbx5* allele^3^. The majority of regions (68%) that were more accessible in the control vs. KO aCM clusters (**Fig. 5b**) were occupied by TBX5 in control aCMs. In contrast, a minority of regions (27%, **Fig. 5b**) with greater accessibility in the KO cluster were TBX5 bound in control aCMs. A similar pattern was found in control vs. knockout early pseudotime clusters (Myocyte_4 vs. Myocyte_6, **Fig. 5b**) – regions with greater accessibility in control were predominantly (53%) TBX5 bound, whereas regions with greater accessibility in knockout were infrequently TBX5 bound (18%). Analysis of patterns of differential accessibility between early vs late and control vs KO clusters showed that the regions could be grouped into those with and those without greatest accessibility in the control aCM cluster. Those with greatest accessibility in the control aCM cluster had functional terms related to cardiac cell development and striated muscle contraction and the majority of these regions were bound by TBX5 (**Supp. Fig. 6**). Together, these data indicate that TBX5 binding is required to maintain accessibility of 1462 regions in control atrial myocytes.

In subsequent analyses, we focused on the regions differentially accessible between the later pseudotime control (Myocyte_1) and KO (Myocyte_6) aCM clusters. There were 1846 regions with significantly greater accessibility in control (“control ATAC regions”) and 356 regions with significantly greater accessibility in *Tbx5^AKO^* (“KO ATAC regions”; **Fig. 5c** and **Supp. Table 5**). We analyzed the contribution of these differentially accessible regions to gene expression. One method of associating genes to regions is by proximity, i.e. pairing each region with its nearest gene. Some of the top differentially accessible regions were nearest to aCM-selective genes *Sbk2*, *Nppa*, *Nppb*, and *Scn5a* (**Fig. 5c**). Other aCM-selective genes neighboring control ATAC regions included *Myl7, Myl4,* and *Bmp10* (**Suppl. Table 5**). *Grid1* and *Smad6* were linked to the most differentially accessible KO ATAC regions (**Fig. 5c**). Grid1 encodes a glutamate receptor related gene that has been shown to be increased in aCMs of humans and dogs in ^36^. *Smad6*, a regulator of TGF and BMP signaling, had greater expression in KO aCMs compared to control (log_2_ fold-change = 1.21, P_adj_ = 1.2E-94). Overall, genes associated with control ATAC regions were more highly expressed in control rather than KO aCMs, whereas genes near KO ATAC regions were more highly expressed in KO aCMs than control (**Fig. 5d**). Genes can also be linked to regions in multiome data by a region-to-gene linkage score based on the nucleus-by-nucleus correlation between the gene’s expression and the region’s accessibility (**Supp. Fig. 7a**).^37^ Repeating the gene expression analysis using this linkage again showed that genes associated with control ATAC regions are more highly expressed than those associated with KO ATAC regions in control aCMs, while those associated with KO ATAC sites were more highly expressed in KO aCMs (**Suppl. Fig. 7b**). These results indicate that control and KO ATAC regions act as cis-regulatory elements that govern expression of neighboring genes.

We next investigated the contribution of control ATAC regions to the regulation of aCM-selective genes. We associated aCM-selective or vCM-selective genes with control and KO ATAC regions, either by proximity or by linkage score. This revealed that 747 (40.4%) control ATAC regions were related to 404 (35.8%) aCM-selective genes (**Supp. Fig. 7c**). In comparison, there were only 243 (13.1%) control ATAC regions related to 42 (4.8%) vCM genes-selective genes. KO ATAC regions also interacted infrequently with aCM-selective (3.6%) or vCM-selective (5.2%) genes, respectively (**Supp. Fig. 7d**). These data suggest that TBX5-dependent accessible regions act as cis-regulatory elements that regulate a substantial subset of aCM-selective gene expression.

To determine the biological significance of the genes associated with control or KO ATAC regions, we performed GO term analysis. Terms enriched for genes associated with control ATAC region involved the regulation of cardiac contraction, development, and conduction (**Fig. 5e**). In contrast, terms enriched for genes associated with KO ATAC regions involved TGFβ and structural terms. These analyses identify regions related to aCM functional properties that are dependent on TBX5 occupancy for accessibility and likely enhancer activity.

To further characterize the differentially accessible chromatin regions and their function as enhancers, we measured their occupancy by the active enhancer marker H3K27ac (**Supp. Fig. 8a-b**). In wild-type aCMs, H3K27ac signal was found at control ATAC regions, and signal was ∼50% lower in *Tbx5^AKO^*. In contrast, wild-type aCMs had relatively weak H3K27ac signal at KO ATAC regions, and signal at these regions was ∼30% greater in *Tbx5^AKO^* aCMs. These data suggest that at least a subset of control and KO ATAC regions act as enhancers in control and *Tbx5^AKO^*, respectively.

We scanned control and KO ATAC regions for known TF binding motifs. In agreement with the high degree of TBX5 occupancy shown by bioChIP-seq, the TBOX motif was the most significantly enriched motif in control ATAC regions, closely followed by the MEF2 motif (**Supp. Fig. 8c**). Footprinting analyses for these motifs demonstrated increased accessibility around the motif, with a sharp loss of accessibility at the motif, consistent with the footprint of TF binding and providing additional support for TF occupancy of these motifs (**Supp. Fig. 8d**). Control ATAC regions were also enriched for other previously identified cardiac TF motifs (**Supp. Fig. 8b**). In contrast, KO ATAC regions were enriched for TF motifs not classically associated with cardiomyocytes, including the NFYA binding site, which was found recently to play a role in CM regeneration after injury (**Supp. Fig. 8e**)^38^.

We further evaluated the binding of cardiac TFs to control and KO ATAC regions using recently reported occupancy data for aCMs and vCMs based on high affinity streptavidin pulldown of endogenously biotinylated alleles of GATA4, MEF2A, MEF2C, NKX2-5, SRF, and TEAD1, in addition to TBX5 and the co-activator P300 (**Supp. Fig. 8f**). At aCM control ATAC regions, TBX5 was the most enriched factor, followed by NKX2-5, which co-binds DNA with TBX5 as a heterodimer^39^. Although the MEF2 motif was the second most enriched motif, its occupancy signal was relatively low compared to the other TFs, with MEF2A being consistently higher than MEF2C. At control ATAC regions in vCMs, TF binding was also observed, although the occupancy signal of all factors was lower than in aCMs. At KO ATAC regions in both aCMs and vCMs, the TF occupancy signal (measured in wild-type aCMs) was lower than in control ATAC regions, TBX5 no longer had the strongest occupancy signal, and no single factor appeared to predominate. Taken together, these data suggest that TBX5 co-occupies control ATAC regions with other key cardiac TFs.

Together, these data show that TBX5 binding positively regulates the accessibility of a large number of atrial enhancers, including enhancers that regulate genes important for aCM identity and function.

### Control ATAC regions participate in TBX5 dependent looping to regulate gene expression

Chromatin is organized into looped conformations that bring enhancer elements into physical proximity of target promoters^40^. To determine if control and KO ATAC regions participate in chromatin loops, we performed H3K27ac HiChIP to identify loops with this active enhancer mark^41^. HiChIP was performed in biological duplicate, each from 10 pairs of AAV9:Nppa-EGFP *Tbx5^Flox/Flox^* or AAV9:Nppa-Cre *Tbx5^AKO^* atria^42, 43^. An average of 375 M read pairs per sample identified 516,995 loops, with 384,293 enhancer-enhancer loops, 122,034 enhancer-promoter loops, and 10,668 promoter-promoter loops (**Fig. 6a; Supp. Table 1**). Statistical comparison identified 510 differential loops between control and *Tbx5^AKO^* (FDR < 0.1), 263 with higher loop score in control (“control loops”), and 247 with higher loop score in *Tbx5^AKO^* (“KO loops”; **Fig. 6b, Supp. Table 6**). Each loop has two loop anchors, and some anchors were shared by more than one loop. As a result, these 263 control and 247 KO loops corresponded to 427 and 128 unique loop anchors, respectively.

**Figure 6.**
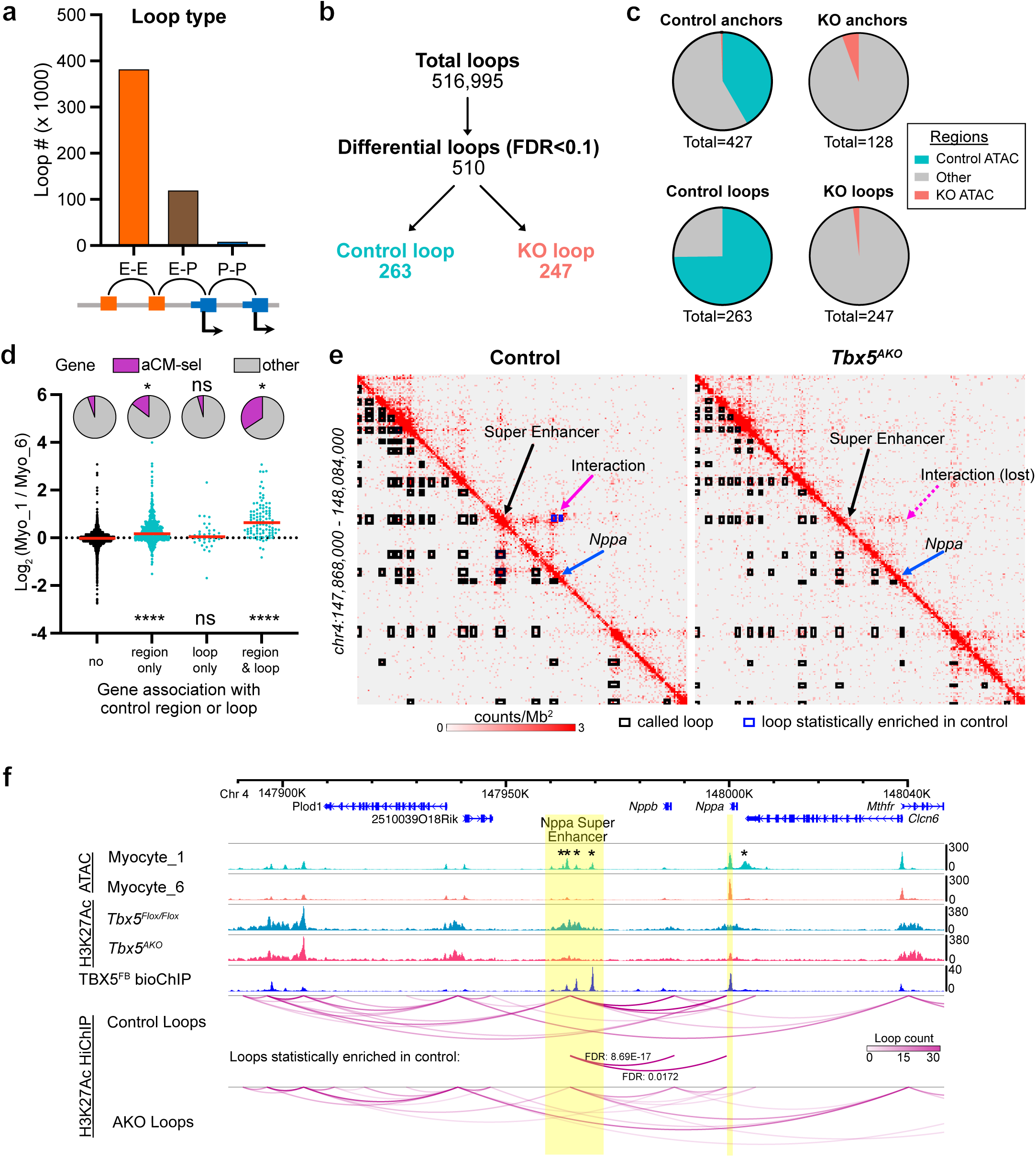
TBX5 maintains local chromatin structure to regulate gene expression. **a,** Loops identified by H3K27ac HiChIP. Loops were classified as enhancer-enhancer, enhancer-promoter, and promoter-promoter. **b**, Differential loops between control and *Tbx5^AKO^* aCMs. Differential looping analysis identified 510 differential loops with FDR<0.1. 263 differential loops had greater loop score in control atrial CMs (“Control loops”) and 247 loops had greater loop score in *Tbx5^AKO^* (“KO loops”). **c**, Overlap of differential loop anchors with control and KO ATAC regions. Top row (anchors) considers each loop anchor individually. Bottom row (loops) considers overlap of at least one anchor from each loop. Control anchors frequently overlapped control ATAC regions, whereas KO anchors rarely overlapped KO ATAC regions and did not overlap control ATAC regions. **d,** Comparison of genes associated with control ATAC region only, control loops only, or both control ATAC regions and control loops. Top: pie charts show the fraction of regions that are aCM-selective. Proportion test vs. “no” association group. Bottom: gene expression ratio in Myocyte_1 vs. Myocyte_6. Kruskal-Wallis vs “no” association group. **e,** Interaction maps for control and *Tbx5^AKO^* samples near the *Nppa* locus. Significant loops are indicated by boxes on the lower left part of the plot. Interactions significantly enriched in control samples are indicated by blue boxes in the upper right-hand region of the plot. *Nppa,* the *Nppa* super enhancer, and their interaction are labeled for control aCMs. The corresponding sites are indicated for *Tbx5^AKO^*aCMs. **f,** Genome browser views of indicated chromatin features. Tbx5^FB^ bioChIP-seq from aCMs is from GEO accession number GSE215065. Yellow highlights indicate *Nppa* super enhancer and Nppa promoter. Asterisks mark regions with significantly enriched accessibility in Myocyte_1 compared with Myocyte_6. Differential looping analysis revealed that connections between the super enhancer and the *Nppa* and *Nppb* gene promoters are present in significantly higher numbers in control samples compared to AKO samples.

To determine how these differences in chromatin looping related to differences in chromatin accessibility, we examined the anchors of control and KO loops for overlap with control and KO ATAC regions. Strikingly, 178 of the 427 (41.7%) unique control anchors, and 196 of 262 (74.8%) control loops overlapped control ATAC regions (**Fig. 6c**). In contrast, only 2 KO loop anchors overlapped control ATAC regions, and only 8 of the 128 (6.3%), and 5 of 247 (2.01%) KO loops, overlapped KO ATAC regions.

We investigated the relationship of chromatin looping to gene expression. By proximity, genes were classified as associated with a control loop without control ATAC region, a control loop with control ATAC region, a control ATAC region that was not part of a control loop, or without association. Genes associated with control loops involving a control ATAC region had the greatest overlap with aCM-selective genes, followed by genes associated with a control ATAC region not part of a control loop (**Fig. 6d**, top). Moreover, the ratio of gene expression between the control and KO aCM clusters was significantly higher in genes associated with control loops and control ATAC regions, followed by genes associated with control ATAC regions but not loops (**Fig. 6d**, bottom). Together, these data support the model that TBX5 is required to maintain enhancer accessibility and local 3D chromatin conformation that supports expression of a subset of aCM genes, including aCM-selective genes. In agreement with this model, most genes near control loops were more highly expressed in control aCM compared to KO aCM clusters and many had aCM-selective expression (**Supp. Fig. 9b**). Genes near KO loops were not enriched for chamber-selective expression and tended to be more highly expressed in KO compared to control aCM clusters (**Supp. Fig. 9c**).

We visualized looping differences at the *Nppa* locus, an aCM-selective gene that was down-regulated in *Tbx5^AKO^* atria (**Fig. 2d** and **Fig. 4e**) and neighbors several control ATAC regions (**Fig. 6f asterisk**). Chromatin contact maps^44^ revealed similar organization of regions flanking this locus in control and *Tbx5^AKO^*. However, at the Nppa locus there was significantly less interaction in *Tbx5^AKO^*of the *Nppa* and *Nppb* promoters with a neighboring, previously characterized super-enhancer that regulates these genes^45^. We visualized this region in a genome browser that included pseudo-bulk ATAC signal from the control and KO aCM clusters, TBX5^FB^ bioChIP-seq, and H3K27ac ChIP-seq and HiChIP loops from control and *Tbx5^AKO^* (**Fig. 6f**)^46–48^. Four control-enriched ATAC regions fell within the *Nppa/Nppb* super-enhancer and were bound by TBX5. The loss in accessibility of these regions in *Tbx5^AKO^* correlated with their significantly reduced participation in loops in *Tbx5^AKO^*. Taken together, these data indicate that TBX5 regulates *Nppa* expression by binding the super-enhancer and promoting its accessibility and linkage to the *Nppa* promoter. *Tbx5* ablation deactivates the super-enhancer in association with *Nppa* down-regulation.

To extend these observations to other critical atrial identity genes, we tested TBX5’s ability to regulate the recently characterized atrial-specific enhancers of *Myl7* and *Bmp10*^3^ (**Fig. 7**). The enhancer regions were cloned into an AAV reporter driving *mCherry* expression and co-injected into neonatal *Tbx5^Flox/Flox^* mice along with either *AAV9:Nppa-EGFP* or *AAV9:Nppa-Cre* (**Fig. 7a).** At P8, hearts were collected and visualized under a fluorescent dissecting microscope. AAV9:Nppa-EGFP treated hearts had EGFP^+^ atria (**Fig. 7b**). The *Myl7* and *Bmp10* enhancers drove *mCherry* expression, which was restricted to the atria. In contrast, pups co-injected with *AAV9:Nppa-Cre* did not have an obvious atrial mCherry signal, indicating that *Tbx5* ablation strongly reduced enhancer activity. To confirm this observation, we analyzed left atrial RNA for *mCherry* transcripts. To control for transduction efficiency, *mCherry* was normalized to expression of a *Broccoli* non-coding RNA driven from U6, an RNA Polymerase III promoter. Expression of *mCherry*, normalized to *Broccoli*, was significantly higher in control compared to *Tbx5*-ablated left atria (**Fig. 7c,d**).

**Figure 7.**
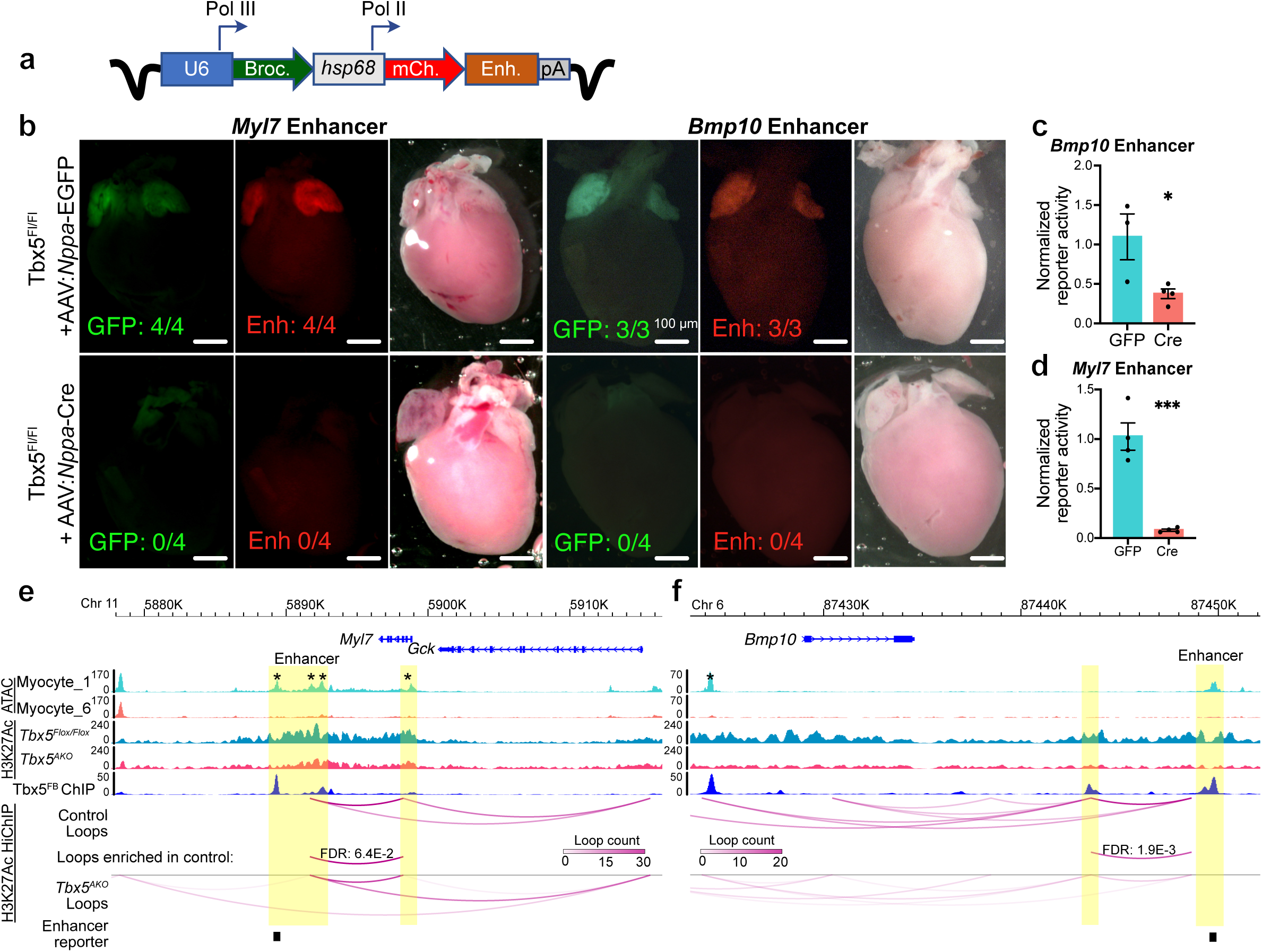
TBX5 promotes atrial gene expression by facilitating the interaction of atrial specific gene promoters with TBX5-dependent enhancers. **a,** AAV enhancer-reporter design. Enhancers of *Myl7* or *Bmp10* (highlighted and labeled in **g** and **h**) were cloned in the 3’UTR of the mCherry reporter gene, expressed downstream of a minimal *hsp68* promoter. The RNA Pol III U6 promoter was used to drive the expression of a *Broccoli* non-coding RNA, which was used to normalize for AAV transduction efficiency. **b,** AAV9 containing either the *Myl7* or *Bmp10* enhancer was co-injected in P2 *Tbx5^Flox/Flox^* with either AAV9:*Nppa-EGFP* (top) or AAV9:*Nppa-Cre* (bottom). Hearts were visualized at P8. Bright field, GFP fluorescent, and mCherry fluorescent images were captured. Number of samples with positive signal and total number of samples is shown. **c-d,** Quantification of mCherry transcripts. *mCherry* transcript levels and *Broccoli* transcript levels were measured by RTqPCR from left atrial RNA. mCherry was normalized to Broccoli to control for transduction efficiency. Error bars represent SEM. Unpaired *t*-test: *, P = 0.0359; ***, P= 0.0005. n=3-4 biological replicates. **e-f,** Coverage plots showing the indicated chromatin features. Tbx5FB bioChip-seq data from aCMs is from GEO accession number GSE215065. Statistically significant differences in looping are diagrammed. Yellow bars indicate *Myl7* enhancer and promoter (e) or *Bmp10* enhancers (f). Asterisks mark regions with significantly reduced accessibility in *Tbx5^AKO^*aCMs.

Chromatin features at these loci illustrated key elements of aCM-selective enhancers. Both of the tested *Myl7* and *Bmp10* enhancers were bound by TBX5 in aCMs (“enhancer reporter” track in **Fig. 7e-f**). The *Myl7* enhancer also contained TBX5-dependent accessible regions (**Fig. 7e**, asterisk) that contacted the Myl7 enhancer via a TBX5-dependent loop (**Fig. 7e**, “Loops enriched in control”). The *Bmp10* enhancer also had greater accessibility in the control compared to the KO aCM cluster, although this did not reach statistical significance (**Fig. 7f**). Furthermore, it contacted a second, TBX5-dependent accessible region through a chromatin loop (**Fig. 7f, asterisk**). The *Bmp10* enhancer also made a TBX5-dependent contact with an additional TBX5 bound region and with the *Bmp10* gene body. Contact maps for the *Myl7* and *Bmp10* regions revealed chromatin contacts anchored in examined enhancer regions to the *Myl7* and *Bmp10* promoters, but with reduced contact in *AKO* samples (**Supp. Fig. 9a**). Together, these data show that TBX5 facilitates promoter-enhancer contacts of atrial genes by binding to and maintaining the accessibility of atrial enhancer elements.

## Discussion

Differences between aCM and vCM gene expression programs imbue them with distinct functional properties required for proper cardiac function. Previously, only one TF, COUP-TFII (also known as NR2F2), was shown to direct specification and differentiation of atrial CMs^12^. However, COUP-TFII regulates early development but not aCM maintenance, since its inactivation later in development did not alter atrial identity^12^. Retinoic acid signaling promotes atrial specification in embryos and in hiPSC-CM differentiation^5, 49–51^. Retinoic acid required NR2F2 and was effective only at early stages of cardiomyocyte specification, indicating that it also influences atrial specification and not maintenance.^5, 50^ Here we demonstrate that TBX5 is necessary to maintain atrial identity in postnatal cardiomyocytes. Postnatal loss of TBX5 downregulated many aCM-selective genes, upregulated vCM-selective genes, and perturbed aCM morphology and function.

Our study identified candidate regulatory regions bound by TBX5 in control aCMs and reduced accessibility in Tbx5^AKO^ aCMs. These regions were decorated by the active enhancer mark H3K27ac, enriched for motifs of TBX5 and other cardiac TFs, most notably MEF2, and co-bound by these TFs. Genes linked to these regions by proximity or by co-variance of accessibility and expression were enriched for aCM-selective expression and for functions important for cardiomyocyte development, structure, and activity. In other words, these represent TBX5-regulated enhancers through which TBX5 influences aCM identity and maintenance of normal cardiac rhythm. This inventory of TBX5-dependent aCM enhancers will be invaluable for future studies of atrial gene regulation in homeostasis and disease.

Enhancers form loops to contact promoters and drive gene expression^52^. H3K27ac HiChIP^41^ measurements revealed changes in enhancer-promoter looping that occurred following *Tbx5* inactivation in aCMs. Remarkably, the large majority of loops that were weakened by *Tbx5* inactivation were anchored on regions that required TBX5 to maintain accessibility. This result suggests that, in addition to maintaining accessibility, TBX5 also maintains the 3D genome structure of a subset of its target enhancers. Since the affected enhancers are enriched for aCM-selective genes including *Nppa*, *Myl7*, and *Bmp10*, this result further implies that TBX5-dependent maintenance of 3D genome structure at key loci is integral to maintenance of the identity of aCMs.

We observed that TBX5 is required to maintain accessibility of thousands of regions. This observation is consistent with recent studies on a human disease-causing *Tbx5* variant, Tbx5-G125R, which increases DNA binding to TBX5 motifs. When this variant was introduced into mice, it increased accessibility of thousands of regions occupied by wild-type TBX5, with associated changes in gene expression and atrial electrophysiology^15^. While TBX5 is not known as a pioneer factor, which binds chromatinized DNA and displaces histones to increase chromatin accessibility, our data indicate that TBX5 binding is required to maintain accessibility. One mechanism by which TBX5 may maintain accessibility is by recruiting chromatin remodeling complexes. TBX5 directly interacts with CHD4 and SMARCD3, subunits of chromatin remodeling complexes, to facilitate heart development^53, 54^. In postnatal aCMs, TBX5 recruitment of these chromatin remodeling complexes may also be required to maintain chromatin accessibility and aCM cardiac gene expression. Further experiments are required to test this hypothesis.

*Tbx5* is essential for maintenance of atrial rhythm^17–19^. Consistent with prior studies of global postnatal *Tbx5* inactivation, and with genome-wide association studies that implicate *TBX5* variants in human AF^55^, our data show that *Tbx5* inactivation selectively in aCMs rapidly results in AF. Prior studies focused on *Tbx5* regulation of Ca^2+^ handling genes as a mechanism underlying AF in *Tbx5*-depleted hearts^18, 19^, which we also observed in our study. More broadly, we observed dysregulation of many atrial identity genes, suggesting that loss of atrial identity and altered expression of these signature atrial genes may also contribute to development of AF. Among signature atrial genes that were most strongly regulated by *Tbx5* was *Sbk2*, a little-studied gene that was recently shown to be essential for aCM sarcomere integrity^33^.

While this study demonstrates an essential role for TBX5 in maintenance of atrial identity of postnatal aCMs, it does not discount other roles of TBX5 in other contexts. Germline inactivation of *Tbx5* disrupts development of both atrial and ventricular chambers^13^, and *Tbx5* is an essential component of the cocktail of TFs that reprogram fibroblasts into cardiomyocyte-like cells^56^. In these contexts, TBX5 is required to promote cardiomyocyte differentiation and proliferation. TBX5 is also expressed in the postnatal vCMs, albeit at lower levels than in aCMs. Overexpression of TBX5 in vCMs was sufficient to promote the expression of atrial genes, suggesting that TBX5 chamber selective function is dose dependent. This dosage sensitivity may contribute to human phenotypes observed with TBX5 variant that increase or decrease expression or activity^15, 20, 57, 58^. Further studies are required to elucidate the molecular mechanisms underlying these dosage- and context-dependent effects of TBX5.

## Supporting information

Supp. Table 1. Summary of high throughput data

Supp. Table 7. Antibodies

Supp. Table 2. DeSeq of TBX5 overexpressing ventricular myocardium

Supp. Table 3. Myo_1 vs Myo_6 DEGS

Supp. Table 4. Differentially accessible regions

Supp. Table 5. Control and KO ATAC regions

Supp. Table 6. Diffloop Hichip

## Author Contributions

MES and YC contributed equally to this study. MES and YC conceived of the study and designed, performed the experiments and analyzed the data. XZ and CP-C. contributed to the data analysis. OB-T, BNA, QM, HW, JMG, MKS, MAT, PW, FL, MG, MP, and RHB contributed data, reagents, and analyses. JGS, CES, IPM, and WTP oversaw the project and provided resources. MES and WTP wrote the manuscript, with input from YC and the other authors.

## Online Methods

### Mice

All research conducted using animals was done following protocols which were approved by Institutional Animal Care and Use Committees at Boston Children’s Hospital, Harvard Medical School, or the University of Chicago. *Tbx5^Flox^*,^13^ Tbx5-OE,^26^ HCN4-CreERT2^59^, Rosa26^mTmG^ ^60^, Rosa26-H2B-mCh,^61^ and Myh6-Cre^62^ mice were described previously. AAVs were injected subcutaneously at the indicated stage.

### Echocardiography

Echocardiography was performed using a Vevo 3100 imaging system (Visual Sonics) with awake animals in a standard hand grip. Cardiac function was recorded from the parasternal short axis view. Atrial size was determined as the ratio of the left atrium to The echocardiographer was blinded to the genotype and treatment of each of the experimental mice.

### Surface EKG

Mice were anesthetized with 3% isoflurane and placed on a heating pad (37°C) for the remainder of the experiment. Mice were positioned in dorsal recumbency and platinum electrodes were placed under the skin of each of the four limbs. 2 minutes of recording were collected, and 1500 beats per mouse were analyzed to calculate the standard deviation of the RR interval or to generate a Poincaré plot. Recordings were performed using Labscribe 3 software, and RR intervals were exported using the same software. SDRR was calculated in Excel. Poincaré plots were generated in R using ggplot2. EKGs were acquired blinded to treatment groups.

### AAV purification

AAVs were purified and quantified following a standard published protocol^63^. Briefly, AAV was produced in HEK293T cells using AAV9 Rep/Cap. Viruses were purified by ionoxidal density gradient ultracentrifugation as described previously^63^. AAV concentration was measured by qPCR. In experiments with HCN4-CreERT2 mice, AAV was injected into the mediastinum via a subxiphoid approach^64^. For all other experiments, AAV (2 x 10^11^ viral genomes per gram bodyweight) was injected subcutaneously to neonatal mice.

### Immunostaining

Tissues were fixed overnight at 4°C, with rotation in 4% PFA. For cryosections, tissues were then placed in 30% sucrose solution until they sank (usually about 4 hours), positioned in OCT and frozen. 5 μm sections were attached to superfrost Plus slides. Sections were permeabilized for 20 minutes using 0.1% triton X, washed, and blocked with 20% donkey serum for 1 hour. Primary antibodies diluted in PBS were incubated on top of tissue sections overnight at 4°C. Slides were washed 3x with PBS and counterstained with the appropriate Alexa-fluor conjugated secondary antibody and Alexa-fluor 647 conjugated WGA (1:500) for 1 hour. Slides were washed 3x, mounted with coverslip solution containing DAPI, and sealed with nail polish. Information about the antibodies used in the study are available in **Supp. Table 6**.

### Microscopy

Confocal images were acquired using a Olympus FV3000RS confocal microscope using a 60x oil immersion lens (NA = 1.4). To measure fibrosis, mouse hearts were excised and fixed in 4% PFA overnight, dehydrated through an ethanol gradient, embedded in paraffin, and sectioned at 7 µm. Sections were dewaxed, rehydrated, stained, and subject to Masson trichrome staining for fibrosis visualization. Sections were imaged by a widefield microscope (Keyence) at ×10 magnification, or a laser scanning confocal (Olympus FV3000RS).

### RTqPCR

Total RNA was isolated using trizol extraction followed by purification using the RNA cleanup and concentrator kit (R1017) and an on-column genomic DNA digest step (Qiagen 79254). Reverse transcription was performed on 500ng total RNA using the Takara Primescript (RR037B). RTqPCR was performed using a Bio-Rad CFX96 or CFX384 real time PCR instrument with Sybr green detection chemistry. Gene expression values were normalized to GAPDH expression. PCR primers sequences: *Myl2-f* CCCTAGGACGAGTGAACGTG, *Myl2-R TCCCGGACATAGTCAGCCTT, Nppa F* GCTTCCAGGCCATATTGGAG, *Nppa R* GGGGGCATGACCTCATCTT, *Nppb F* GAGGTCACTCCTATCCTCTGG, *Nppb R* GCCATTTCCTCCGACTTTTCTC, *Myl7 F* GGCACAACGTGGCTCTTCTAA, *Myl7 R* TGCAGATGATCCCATCCCTGT, *mCherry F* ACGGCCACGAGTTTGAGATT, *mCherry R* CAAGTAGTCGGGGATGTCGG, *Broccoli F* TATTCGTATCTGTCGAGTAGAGT, *Broccoli R* GATCATCAGAGTATGTGGGAG*, GAPDH F* AGGTCGGTGTGAACGGATTTG, *GAPDH R* TGTAGACCATGTAGTTGAGGTCA. *Tbx5 F* GGCATGGAAGGAATCAAGGT, *Tbx5 R* CTAGGAAACATTCTCCTCCCTGC.

### Western Blotting

Atrial and ventricular lysates were prepared by homogenizing samples in ice-cold RIPA buffer containing protease inhibitor. Protein concentrations for the different samples were determined using pierce BCA Protein Assay Kit from Thermofisher. An equal amount of protein (20 µg) was loaded into each well of an SDS-PAGE gel. Gels were transferred to PVDF membranes, which were blocked with 5% milk, and incubated with primary antibodies overnight at 4°C (Supp. Table 3). Secondary antibodies conjugated to HRP were incubated the following day for 1 hr. The membranes were then developed and imaged using the ImageQuant LAS 4000 Luminescent Image Analyzer.

### SnRNA and snATAC sequencing

Multiome samples were flash frozen in liquid nitrogen and preserved at −80 degrees celsius prior to the experiment. To minimize experimental differences, all samples were processed on the same day and Gel Bead-In EMulsions were encapsulated in the same run. Nuclei were isolated following a published protocol with minor modifications^65^. Briefly, whole frozen atria were resuspended in homogenization buffer (250 mM sucrose, 25 mM KCl, 5 mM MgCl_2_, 10 mM Tris, pH 8, 1 µM DTT, with added protease inhibitor, 0.4 U/µl RNaseIn, 0.2 U/µl SuperaseIn, and 0.1% Triton X-100). Samples were then homogenized in a Qiagen TissueLyser II with a 1 mm ball bearing set at 25 hz for 90 seconds. Samples were filtered using a pluriselect ministrainer (40 µm), centrifuged for 5 minutes at 500g, and resuspended in storage buffer (4% BSA in PBS, 0.2 U/µl RNaseIn). Samples were stained using 7-aminoactinomycin D (7-AAD) at a final concentration of 1 µg/ml, and FACS sorted for purification. For each multiome sample, 300,000 nuclei were sorted into storage buffer. Nuclei were then gently permeabilized using multiome lysis buffer (Tris-HCl Ph 7.4 10 mM, NaCl 10 mM, MgCl_2_ 3 mM, Tween-20 0.1%, NP40 0.1%, Digitonin 0.01%, BSA 1%, DTT 1mM, RNaseIN 1 U/µl), washed 2x with wash buffer (Tris-HCl pH 7.4 10 mM, NaCl 10 mM, MgCl_2_ 3 mM, BSA 1%, Tween-20 0.1%, DTT 1 mM, RNaseIN 1 U/µl), and resuspended in 1x nuclei buffer (10x Genomics). Nuclei were quantified using the Countess 3 system, and then processed using the 10x Genomics Next GEM Single cell multiome ATAC + Gene Expression Reagents kit. The resulting joint snRNA and snATAC libraries were analyzed by tapestation and then sequenced at the depth recommended by 10x Genomics.

### Multiome analysis

Outputs from the 10x Cellranger-ARC software package were processed using the Seurat^29^ and Signac^27^ packages. Briefly, Cellranger-ARC was ran for each sample, and the output files were used to construct Seurat objects in R. A shared peak set derived from the integration of peaks identified for each sample was used to analyze the snATAC dataset, according to the merging Signac vignette. Each Seurat object was QC’d using the same metrics: 250 < nCount_ATAC < 100,000; 250 < nCount_RNA < 35,000; nucleosome_signal < 2; TSS.enrichment > 1; and percent mitochondrial reads < 25. Next, doublets were identified and removed using Doubletfinder^66^ on the RNA portion of the assay. Finally, the objects were merged together. Pre-processing and dimensional reduction were performed on the RNA and ATAC assays independently, using standard approaches for RNA and ATAC-seq data^67^, as follows. Gene expression UMI count data was normalized using SCTransform, and the 20 nearest neighbors (KNN, *k* = 20) for each nucleus was found using the FindNeighbors function. LSI was used to perform dimension reduction on the DNA accessibility assay dataset, and graph-based clustering on the LSI components 2:15 was performed by computing a nearest-neighbor graph using LSI low-dimensional space (*k* = 20 neighbors) and then applying the Smart Local Moving algorithm for community detection. A WNN graph was generated using the weighted combination of the RNA and ATAC modalities. All downstream analysis were performed using features of Seurat/signac or using Seurat wrappers for other packages. Trajectory analysis was performed using the Seurat wrapper for Monocle3^30^. Differential gene expression and differential genomic accessibility analyses were performed comparing clusters using the FindMarkers function on either the RNA or the ATAC assays. Differential accessibility was performed using logistic regression with the total number of ATAC fragments as a latent variable, with thresholds of | log_2_FC | > 0.5 and P_Adj_ <0.05. The same thresholds were used to identify differentially expressed genes. Differential gene expression analysis was performed using the Wilcoxon test. To generate bigwig files from the ATAC data for individual clusters, fragment files for each of the samples were split using the list of cell barcodes composing the cluster of interest using the sinto package into .bam files. The bamcoverage tool was used to create normalized .bigwig files (RPGC normalization).

### Gene set analysis

We defined aCM-selective and vCM-selective genes based on bulk RNA-seq of P0 aCMs vs P0 aCMs^3^. Each set of selective genes was defined by log_2_ fold change > 2, padj<0.05, and TPM≥ 5. Skewd distribution of a setog genes among a list of genes rank ordered by relative expression intwo conditions was analysed using Gene Set Enrishment Analysis^68^. Enrichment of a list of genes in predefined gene pathways (“GO term enrichment analysis”) was performed using the DE and enrichR pathway visualization tool through Seurat, with default settings^69^. Enrichment of genes linked to regions in predefined gene pathways was performed using GREAT^70^.

### Bulk RNA-seq

Paired -end reads were aligned to the mm10 genome with STAR^71^ version 2.5.3b using the parameter —quantMode GeneCounts to retrieve counts at exons. Genes were tested for differential expression with DESeq2 with default settings. Genes differentially expressed between Tbx5-OE and control ventricles were those with Padj < 0.05 and | log2 fold-change | > 0.5.

### H3K27ac HiChIP

HiChIP for H3K27ac was performed using the Arima-HiC+ kit for HiChIP and the library was prepared using the Swift Biosciences Accel-NGS 2S Plus DNA Library Kit following the manufacturer’s protocol. HiChIP samples (10 left and right atria per biological replicate) were collected and snap-frozen in liquid nitrogen. Nuclei were isolated using the same approach as the multiomics experiment. After homogenization in the TissueLyser II, nuclei were pelleted and resuspended in PBS. Nuclei from aCMs were enriched by purification by MACS for PCM1, as previously described^72^. Formaldehyde was added to the nuclei for a final concentration of 2%, and the nuclei were fixed with gentle rocking at room temperature for 15 minutes. The reaction was quenched with glycine, and the nuclei were pelleted and counted. In total, each HiChIP reaction required ∼3 million nuclei. The fixed nuclei were pelleted and snap frozen and stored at −80°C until used in the HiChIP protocol. Libraries resulting from the HiChIP protocol were sequenced to a depth of 300 million reads per library.

H3K27ac ChIP signal was obtained by ChIP-seq analysis of the HiChIP data.

### Loop calling and differential loop identification

FASTQ files from the HiChIP experiment were used to generate normalized contact matrices with HIC-PRO^42^. Then, the command line tool hichipper^43^, a restriction enzyme site aware package that reduces the bias caused by the digest step of HiChIP, was used to identify loop anchors and to call loops, producing .mango interaction files that contain information on all significant contacts and anchor regions^73^. To determine if loops were significantly enriched in control or KO samples, interaction .mango files for each biological replicate were loaded into the R package diffloop^74^. To determine significance, the default edgeR statistical test used by diffloop was applied, where counts are modeled with negative binomial distribution and the empirical Bayes procedure is used to mediate overdispersion. Loops that were called in one biological replicate greater than 5 times but not the other and interactions < 5 kb in length were excluded. Significant loop counts were multiplied by a size factor to normalize for sequencing depth, and mean values less than 5 were filtered out. Interaction types were determined by annotating the genome. Regions (+/− 1 kb) from the TSS of mm10 genes were labeled as promoters, and other peaks called by MACS2 for each of the samples IP’d for the enhancer mark H3K27ac were defined as enhancer regions.

### Generating contact maps of HiChIP data

To construct contact maps and .bigwig files, FASTQ reads were aligned to the mm10 genome using BWA^75^, and ligation junctions were identified with pairtools^76^. PCR duplicates were removed and .bam and .pairs files were then generated with pairtools. The resulting mapped.pairs file was used as the input to generate a .hic file for each sample with juicer tools. Juicebox was used to visualize contact matrices^77^. Loops were identified on the resulting contact matrices by loading the .mango file output from hichipper from a given sample into Juicebox. Resulting .bam files were used as input with the bamCoverage tool to generate bigwig files, which were normalized using ‘RPGC’ to the effective mm10 genome size. Heatmaps and profiles were generated from .bam files of control and KO ATAC regions using deeptools.

### Data Access

High throughput data used in this manuscript are available from the Gene Expression Omnibus, accession number (pending). Other datasets used were obtained from GSE215065^3^ and GSE129503^17^.

### Statistics

Results were expressed as mean ± SEM. Statistical tests, indicated in figure legends, were performed using Graphpad Prism 9 or R. P<0.05 was used as the statistical threshold for significance.

