## Supplemental Figures for "*Tbx5* maintains atrial identity by regulating an atrial enhancer network"

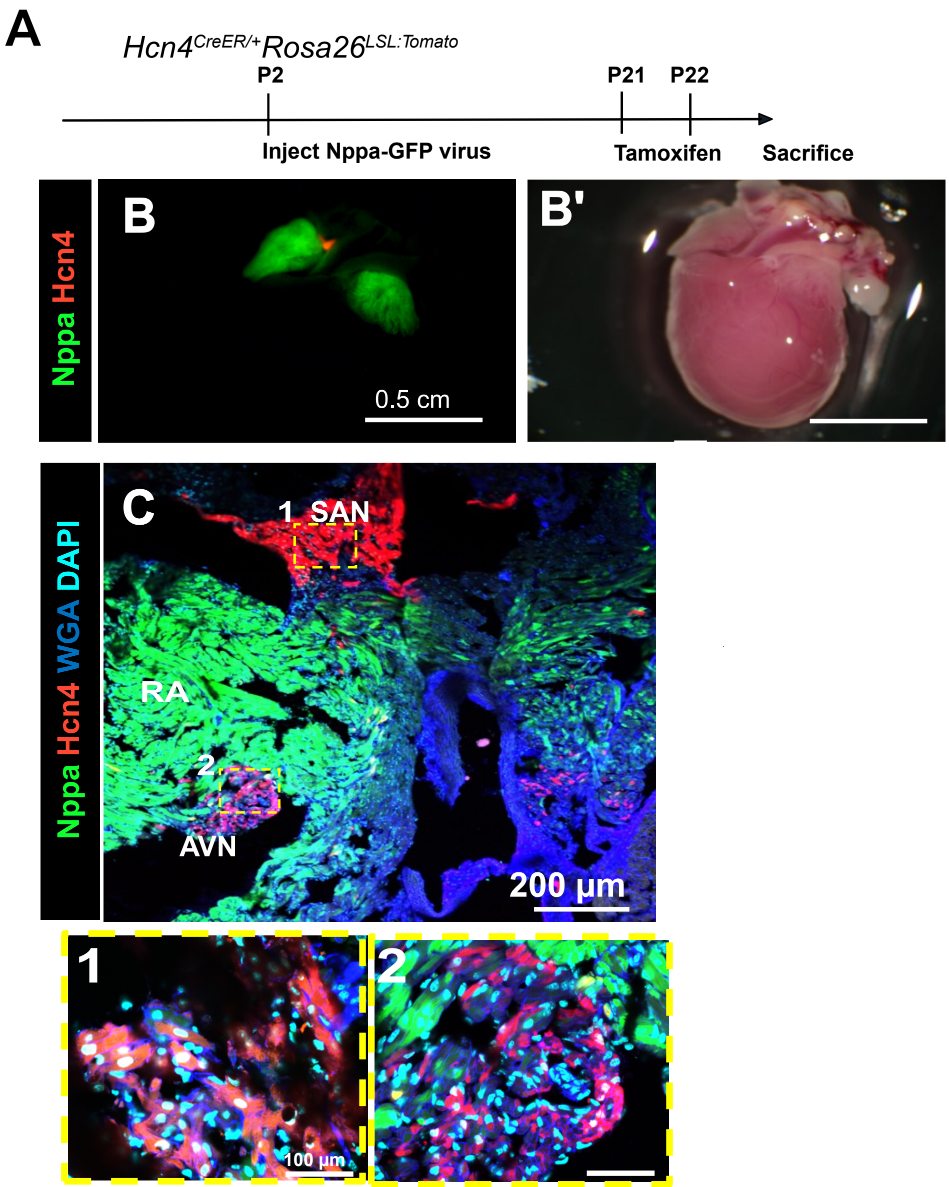


**Supplemental Figure 1. Characterization of the cardiac expression domain of AAV9:Nppa-EGFP. a-b,** Hcn4$^{CreERT2/+}$; Rosa26$^{LSL\text{-}Tomato}$ pups were injected with 2 X $10^{11}$ viral genomes per gram bodyweight (VG/g) AAV9:Nppa-EGFP at P8 and fed tamoxifen at P21 and P22. Hearts were harvested in PBS and brightfield, GFP, and Tomato fluorescent images were acquired. **c**, Hearts were sectioned and stained with WGA. Tomato signal marked the sinoatrial node (SAN) and atrioventricular node (AVN) but not the working myocardium. GFP was restricted to the working myocardium.

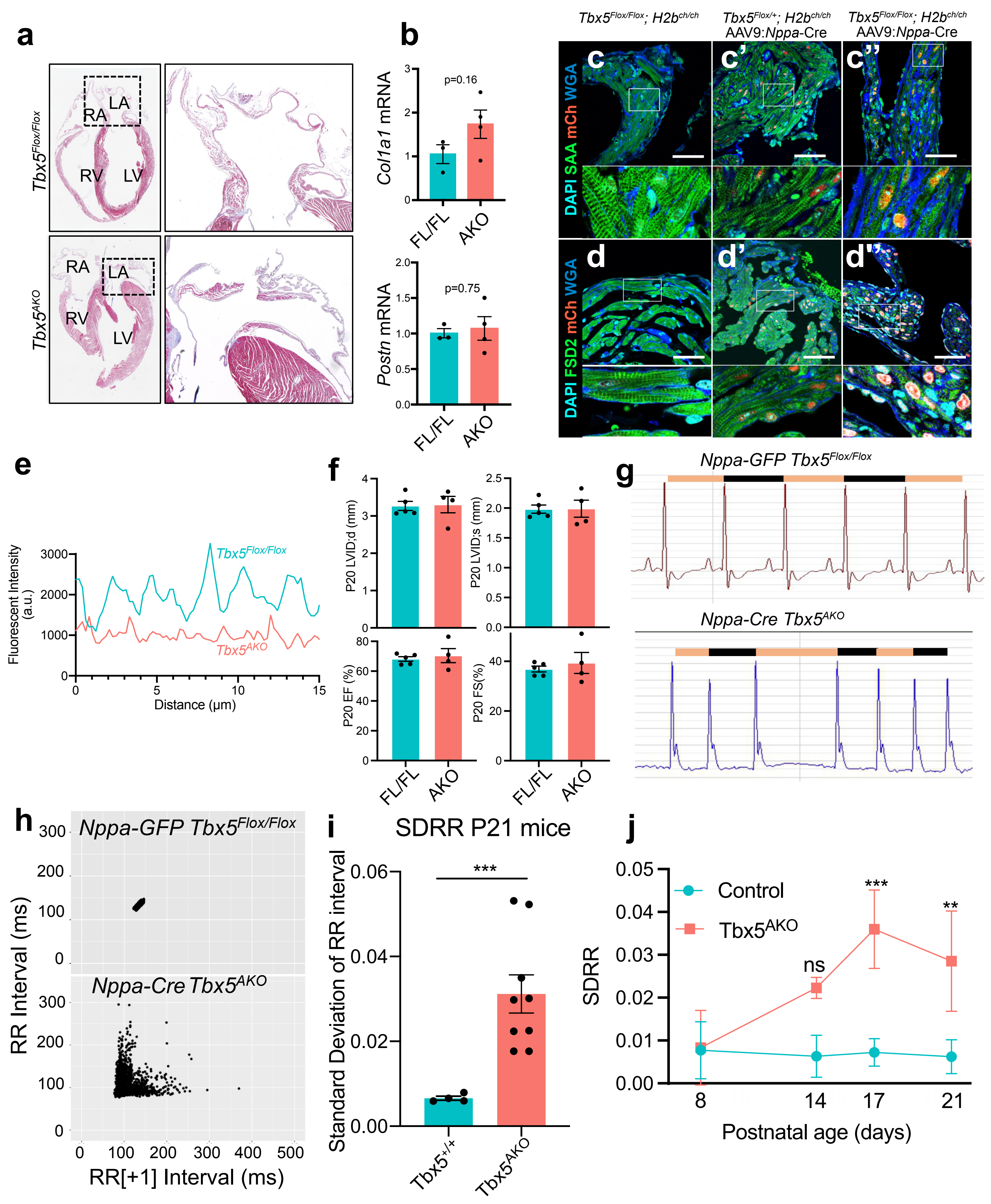


**Supplemental Figure 2. Phenotypic characterization of Tbx5$^{AKO}$ mice. a**, Trichrome staining of Tbx5$^{Flox/Flox}$ and Tbx5$^{AKO}$ hearts. **b,** RTqPCR quantification of *Postn* (periostin) and *Col1a1* (collagen type 1 alpha 1 chain), two indicators of fibrosis. **c-d**, Immunostaining for sarcomeric α-actinin (SAA) or FSD2, markers of the Z line and the junctional Sarcoplasmic reticulum, respectively, in the left atrium of the indicated genotypes. Localization of both proteins is disrupted in Tbx5$^{AKO}$ atria. **e**, Pattern (continued)

**Suppl. Fig. 2, continued.**

of SAA signal intensity. SAA intensity along the long axis of cardiomyocytes demonstrated a periodic signal in control, consistent with regular position of sarcomere Z-lines, and loss of periodicity in $Tbx5^{AKO}$. **f**, Preserved ventricular function of $Tbx5^{AKO}$ mice. $Tbx5^{Flox/Flox}$ mice were treated with AAV9:Nppa-EGFP (control) or AAV:Nppa-Cre ($Tbx5^{AKO}$) at P2. Echocardiography was performed at P20. LVID;d, left ventricular internal diameter at end diastole. EF, ejection fraction. **g**, Surface EKG recordings. Orange and black bars highlight successive RR intervals. **h,** Poincaré plots. The RR interval of greater than 1500 beats on EKG recordings is plotted versus the RR interval of the subsequent beat (RR[+1]). to visualize the dispersion of interbeat intervals in $Tbx5^{AKO}$, consistent with atrial fibrillation. **i**, Standard deviation of the RR interval, a measure of heart rate irregularity, was calculated for at least 1500 beats for each group at P21. Unpaired t-test: ***, P<0.001. **j**, Time course of heart rate irregularity. Serial EKGs were acquired from control or $Tbx5^{AKO}$ mice at the indicated time points. SDRR was measured and compared between groups using a two-way ANOVA. Sidek's multiple comparison test was used to compare between genotypes at each time point. **, P<0.01. ***, P<0.001.

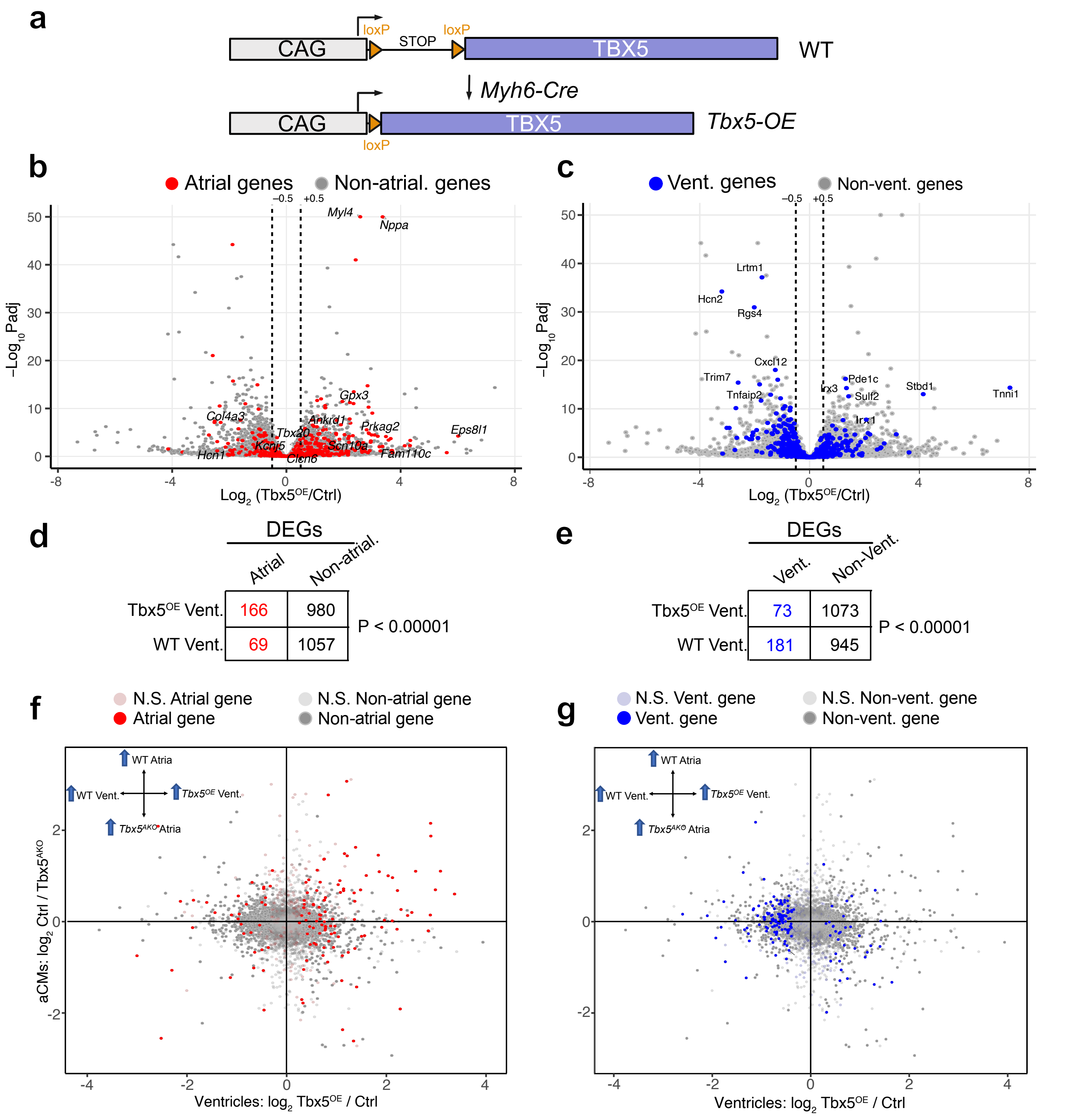


**Supplemental Figure 3. *Tbx5* overexpression atrializes ventricular myocytes. a**, Strategy to generate TBX5-OE ventricles. A full length *Tbx5* cDNA downstream of the ubiquitous CAG promoter is activated by the cardiomyocyte specific Myh6-Cre transgene. **b-c**, Volcano plot comparing the change in gene expression of mouse left ventricle overexpressing *Tbx5* compared to control LV. aCM genes are marked in red in (a) and vCM genes are denoted in blue in (b). **d-e**, Fisher's exact test was performed to determine if changes in chamber selective gene expression downstream of *Tbx5* overexpression were significant. **f-g**, Combining the effect of Tbx5 knockout in atria and Tbx5 overexpression in ventricles. aCM genes are denoted in red (f) and vCM genes are denoted in blue (g). Many of the genes dependent on *Tbx5* in atria and upregulated by *Tbx5* in ventricles (upper right quadrant) are aCM genes. vCM genes tended to be downregulated by *Tbx5* overexpression in ventricles.

**a**

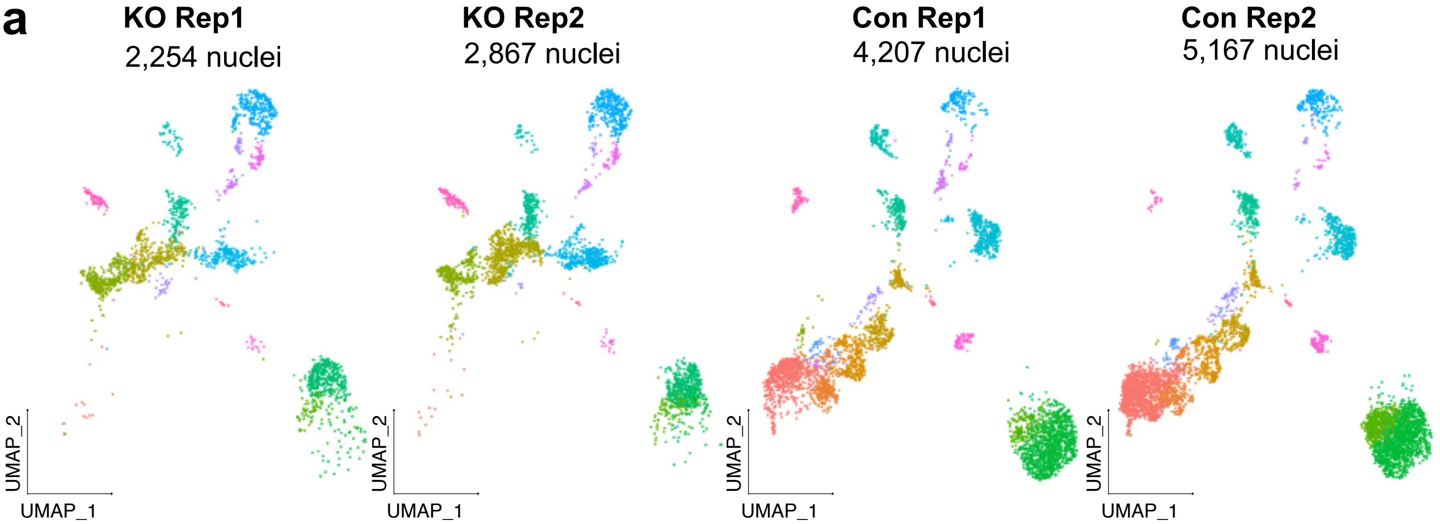


**b**

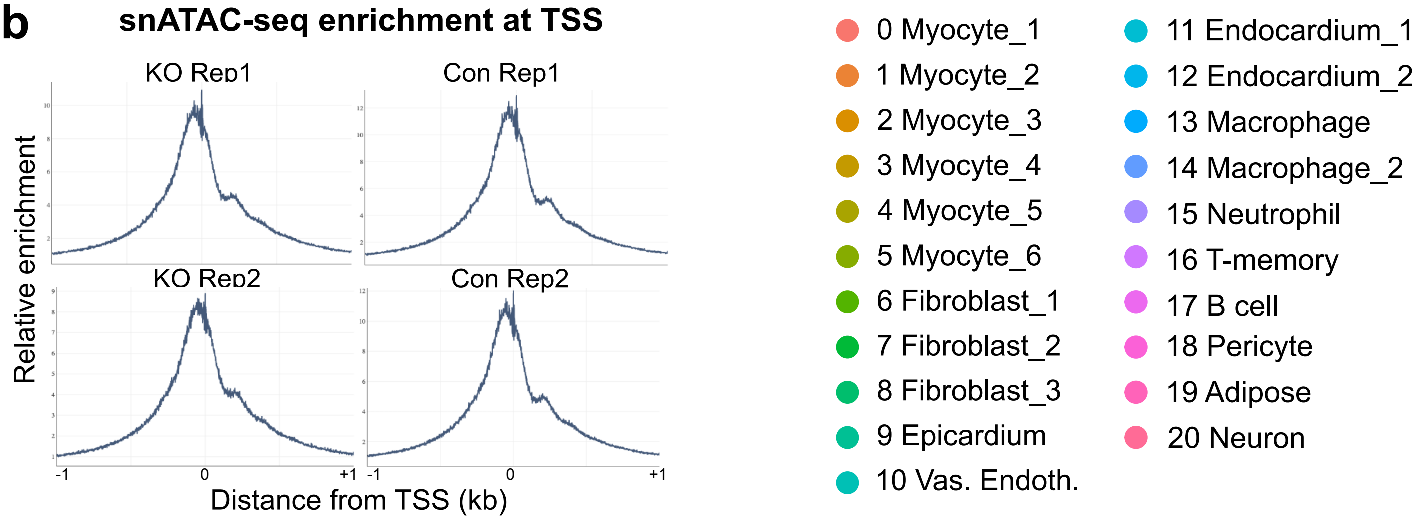


**c**

| Sample | mean nCount_ATAC | mean nFeature_ATAC | mean nCount_RNA | mean nFeature_RNA |
|---|---|---|---|---|
| KO1 | 10778.79 | 7880.126 | 4892.911 | 1969.715 |
| KO2 | 9809.963 | 7084.746 | 4760.472 | 1913.099 |
| Con1 | 7559.528 | 5823.415 | 6431.626 | 2153.253 |
| Con2 | 6717.072 | 5190.006 | 4504.234 | 1731.528 |

**Supplemental Figure 4. Single cell dataset metrics. a,** WNN UMAP of the dataset split by original sample. Each of the KO and control replicates have a high degree of overlap, demonstrating high reproducibility of cell state changes in Tbx5$^{AKO}$ atria. **b**, Transcription start site (TSS) aggregation plots for the scATAC multiome datasets showing the expected enrichment of ATAC fragments. **c**, Parameters from single nucleus datasets. Mean values per nucleus: nCount_ATAC, number of ATAC fragments; nFeature_ATAC, number of ATAC peaks with at least one read; nCount_RNA, number of RNA fragments; and nFeature_RNA, number of genes.

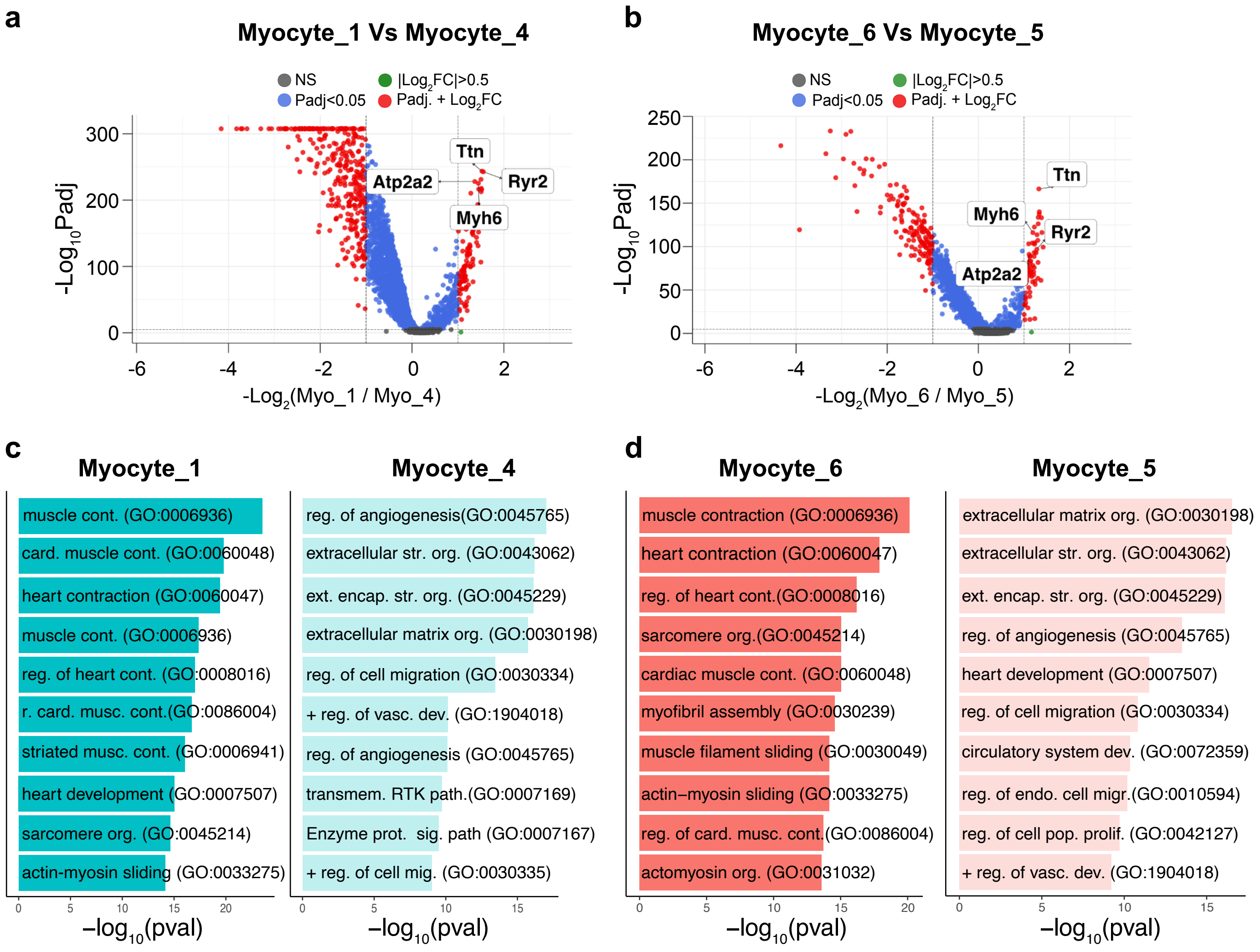


**Supplemental Figure 5. Differentially expressed genes between myocyte clusters. a**, Volcano plot of differentially expressed genes (DEGs) between the control clusters Myocyte_4 (left) and Myocyte_1 (right). **b**, Volcano plot of the DEGs between the KO clusters Myocyte_5 (left) and Myocyte_6 (right). **c-d**, GO biological process terms enriched for DEGs for the comparisons shown in (a) and (b). The top 10 terms are shown. to compare the different myocyte terms. Myocyte_1 and Myocyte_6 are enriched for terms related to cardiac function, demonstrating the relative maturity of these CM populations compared to Myocyte_4 and Myocyte_5, which are enriched for developmental and remodeling terms.

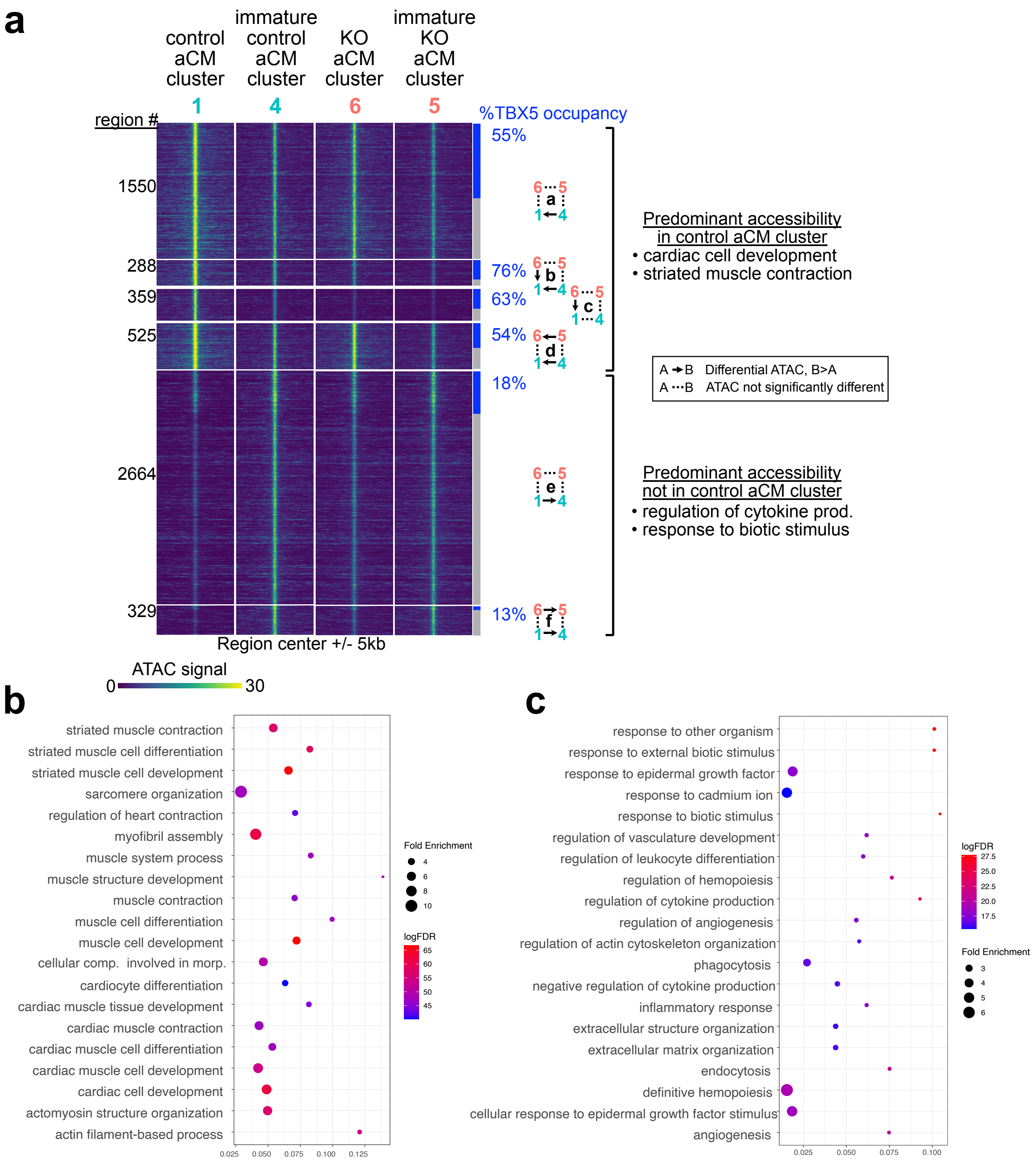


**Supp. Figure 6. Characterization of myocyte differentially accessible regions.** Heatmap (**a**) shows the patterns of differential accessibility between myocyte clusters. The rows contain the union of regions with differential accessibility in the four pairwise comparisons shown in Fig. 5a. ATAC signal in each region is shown for myocyte clusters 1 (control aCM cluster), 4, 6 (KO aCM cluster), and 5. The regions are grouped (groups a-f) by their pattern of accessibility change in the four pairwise comparisons. Arrows denote significant enrichment in one cluster compared to another. These six groups fit two predominant patterns: those with and those without predominant accessibility in the control aCM cluster (Myocyte_1). Most regions with predominant accessibility in the control aCM cluster were occupied by TBX5 and had GO terms related to cardiac cell development or striated muscle contraction (**b**). In contrast, a minority of regions without predominant accessibility in the control aCM cluster were occupied by TBX5 and had GO terms that were atypical for cardiomyocytes (**c**). Groups with less than 200 regions are not shown in the heatmap.

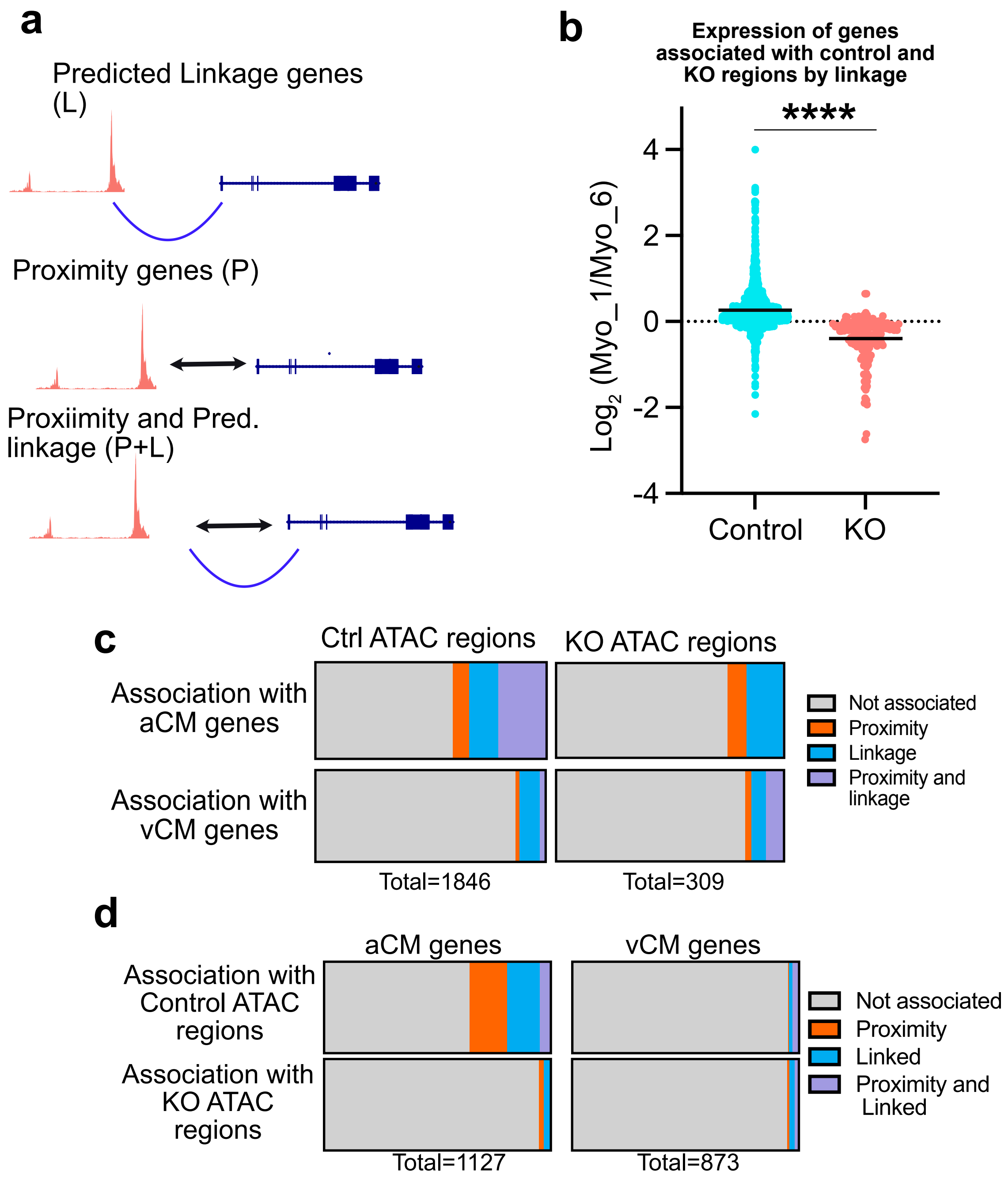


**Supplemental Figure 7. Proximity and predicted gene linkages demonstrate the regulation of the atrial GRN by control peaks. a**, The different types of association of a genomic region with a gene: (1) Linkage (L). Region-to-gene linkages are predicted based on co-variance of accessibility and expression on a nucleus-by-nucleus basis in the multiome data. (2) Proximity (P). Region-to-gene relationships are inferred by proximity of the region to the gene's transcriptional start site. (3) Proximity and linkage (P+L). A region-to-gene association can be supported by both proximity and linkage. **b,** Genes were associated with control and KO regions by linkage. Ratio of gene expression between the control and KO aCM clusters was plotted and compared between groups by the Mann-Whitney test. ****, P<0.0001. **c**, Association of control and KO ATAC regions with aCM and vCM genes. Associations were made on the basis of proximity, linkage, or both. **d**, aCM and vCM genes were interrogated for associations with control and KO ATAC regions by proximity, linkage, or both.

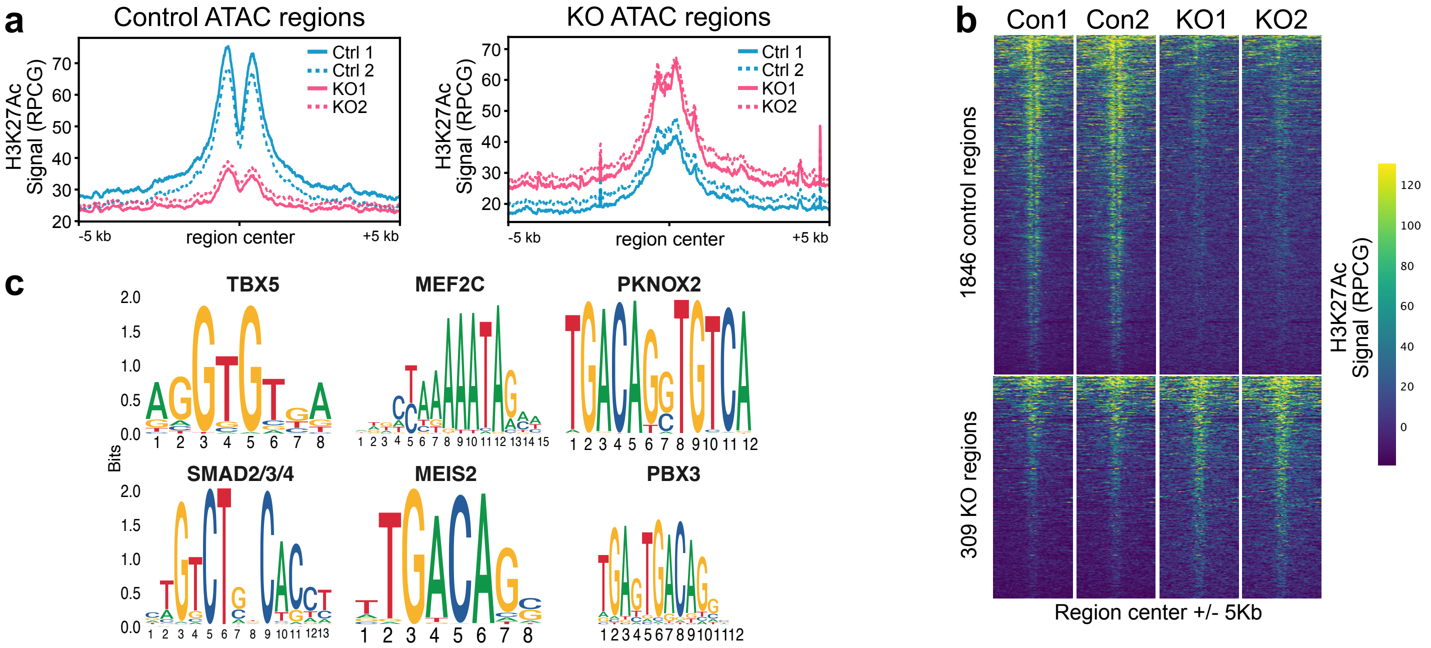


| motif | percent.observed | percent.background | fold.enrichment | pval_Adj | motif.name |
|---|---|---|---|---|---|
| MA0807.1 | 25.5 | 9.7 | 2.6 | 5.88E-51 | TBX5 |
| MA0497.1 | 36.6 | 25.1 | 1.5 | 5.09E-17 | MEF2C |
| MA0783.1 | 23.7 | 14.5 | 1.6 | 5.07E-16 | PKNOX2 |
| MA0513.1 | 33.7 | 23.1 | 1.5 | 1.91E-15 | SMAD2/3/4 |
| MA0797.1 | 19.4 | 11.4 | 1.7 | 7.49E-15 | TGIF2 |
| MA0774.1 | 17.5 | 10.7 | 1.7 | 7.15E-12 | MEIS2 |
| MA0782.2 | 24.7 | 17.0 | 1.5 | 8.49E-11 | PKNOX1 |
| MA1114.1 | 26.6 | 19.2 | 1.4 | 2.27E-09 | PBX3 |
| MA1576.1 | 21.8 | 15.7 | 1.4 | 6.79E-08 | THRB(var.3) |
| MA0595.1 | 14.9 | 10.0 | 1.5 | 3.03E-07 | SREBF1 |
| MA0093.3 | 17.6 | 12.4 | 1.4 | 3.60E-07 | USF1 |
| MA0831.2 | 12.4 | 8.0 | 1.6 | 4.30E-07 | TFE3 |
| MA1494.1 | 15.7 | 10.8 | 1.5 | 4.41E-07 | HNF4A(var.2) |
| MA1485.1 | 17.0 | 11.8 | 1.4 | 4.45E-07 | FERD3L |
| MA0526.3 | 17.0 | 12.0 | 1.4 | 1.37E-06 | USF2 |
| MA0730.1 | 18.3 | 13.2 | 1.4 | 1.49E-06 | RARA(var.2) |
| MA1467.1 | 24.8 | 19.1 | 1.3 | 2.37E-06 | ATOH1(var.2) |
| MA0596.1 | 15.3 | 10.7 | 1.4 | 3.18E-06 | SREBF2 |
| MA0504.1 | 26.7 | 21.0 | 1.3 | 5.23E-06 | NR2C2 |
| MA0499.2 | 22.5 | 17.3 | 1.3 | 7.35E-06 | MYOD1 |
| MA0672.1 | 13.6 | 9.7 | 1.4 | 1.75E-05 | NKX2-3 |

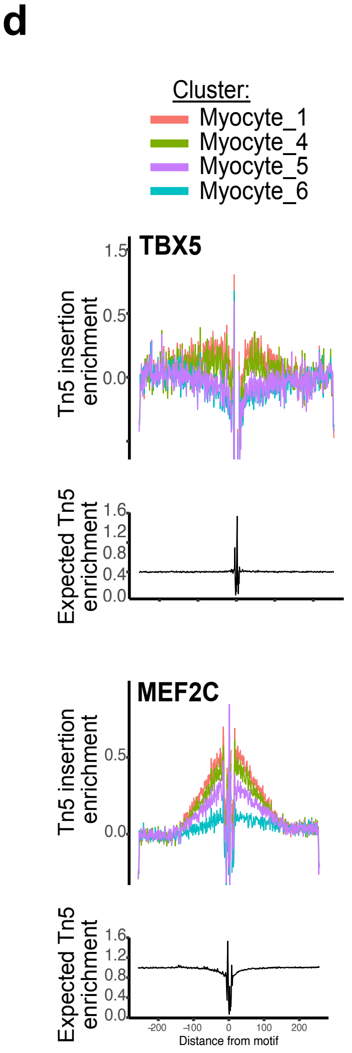


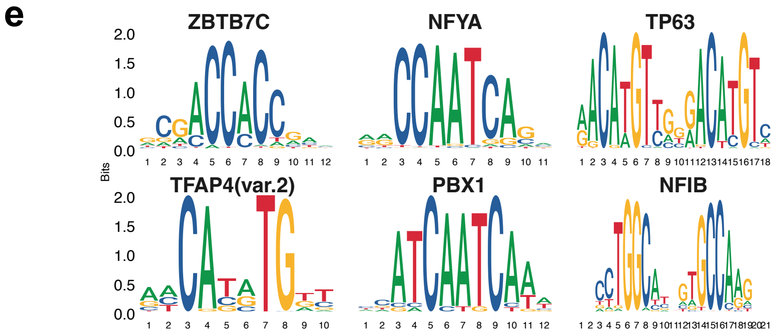


| motif | percent.observed | percent.background | fold.enrichment | pval_Adj | motif.name |
|---|---|---|---|---|---|
| MA0695.1 | 15.1 | 5.1 | 3.0 | 1.89E-20 | ZBTB7C |
| MA0060.3 | 19.1 | 8.6 | 2.2 | 4.22E-16 | NFYA |
| MA0525.2 | 12.9 | 4.9 | 2.6 | 1.12E-14 | TP63 |
| MA1570.1 | 14.4 | 6.3 | 2.3 | 1.02E-12 | TFAP4(var.2) |
| MA0070.1 | 15.4 | 7.1 | 2.2 | 2.13E-12 | PBX1 |
| MA1643.1 | 32.0 | 21.3 | 1.5 | 6.79E-10 | NFIB |
| MA1117.1 | 16.4 | 9.0 | 1.8 | 4.98E-09 | RELB |
| MA0747.1 | 38.4 | 27.6 | 1.4 | 5.18E-09 | SP8 |
| MA0056.2 | 19.9 | 12.1 | 1.6 | 2.54E-08 | MZF1 |
| MA0073.1 | 55.6 | 44.8 | 1.2 | 5.38E-08 | RREB1 |
| MA1653.1 | 48.4 | 37.9 | 1.3 | 8.56E-08 | ZNF148 |
| MA1523.1 | 17.8 | 10.7 | 1.7 | 1.21E-07 | MSANTD3 |
| MA0741.1 | 43.9 | 33.8 | 1.3 | 1.49E-07 | KLF16 |
| MA1581.1 | 25.8 | 17.5 | 1.5 | 1.98E-07 | ZBTB6 |
| MA0143.4 | 16.8 | 10.2 | 1.6 | 5.50E-07 | SOX2 |
| MA1592.1 | 19.4 | 12.4 | 1.6 | 5.61E-07 | ZNF274 |
| MA1120.1 | 17.1 | 10.6 | 1.6 | 8.45E-07 | SOX13 |
| MA1522.1 | 37.7 | 28.8 | 1.3 | 1.43E-06 | MAZ |
| MA1528.1 | 23.5 | 16.2 | 1.4 | 2.37E-06 | NFIX(var.2) |
| MA0695.1 | 15.1 | 5.1 | 3.0 | 1.89E-20 | ZBTB7C |

**Supplementary Figure 8. Motif analysis of control and KO ATAC regions.**

**f**

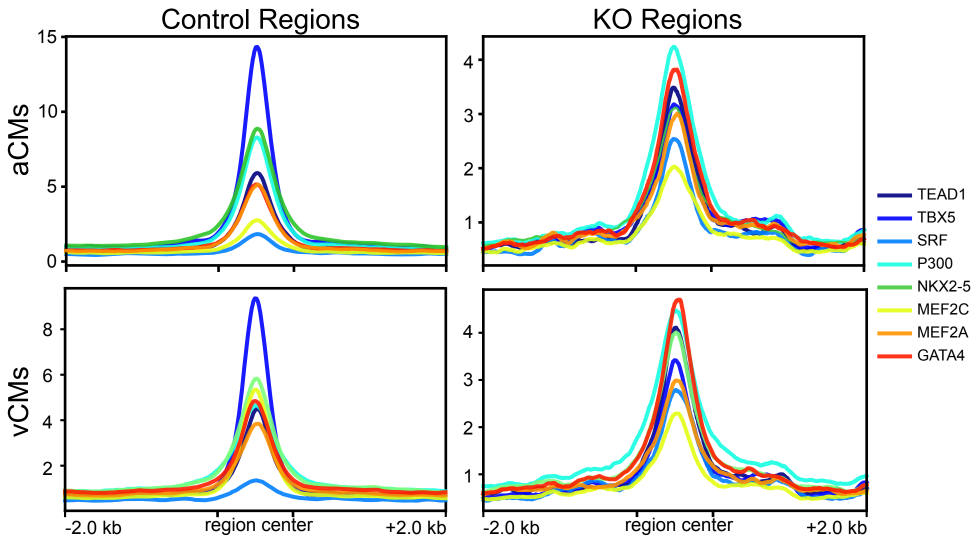


**Suppl. Fig. 8, continued.**

**a**, Aggregation plots for H3K27Ac at control and KO regions. **b**, Heatmaps of the H3K27Ac signal at control and KO peaks. **c**, Top-enriched motifs identified in control regions. The TBOX motif was the most enriched, followed by MEF2. An extended table of non-redundant motifs showing the top 21 most enriched motifs in control regions. **d**, Transcription factor footprinting analysis demonstrates footprints at Tbox and Mef2 motifs in control clusters (Myocyte_1 and Myocyte_4) compared to KO clusters (Myocyte_5 and Myocyte_6). **e**, Top-enriched motifs identified in KO regions and extended table of the top non-redundant motifs. **f**. Occupany of control and KO ATAC regions by cardiac TFs in aCMs. Occupancy data is from GEO GSE215065[24].

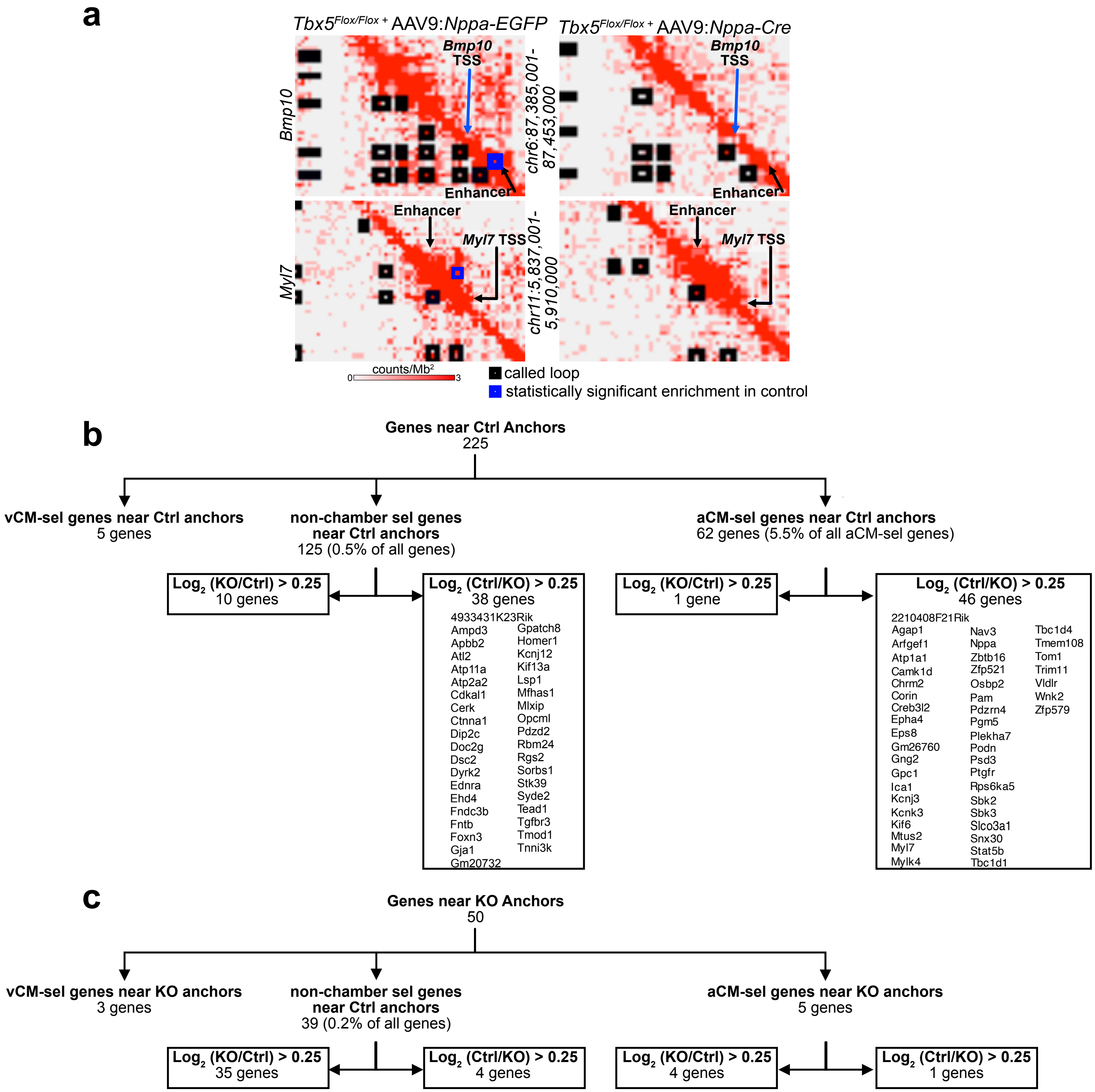


**Supplemental Figure 9. Chromatin loops link TBX5 dependent enhancers with atrial genes. a**, Contact maps of *Myl7* or *Bmp10*. Black boxed regions are loops called in each sample and blue boxed regions mark differential loops that are significantly stronger in control samples. **b**, Genes near control anchors grouped by adjacency to aCM-selective, vCM-selective, or non-chamber selective expression. Control anchors were present near 62 aCM-selective genes and of these, 46 were expressed at greater levels in control Myocyte_1 compared to KO Myocyte_6. Notable genes from previous figures include *Nppa*, *Bmp10*, *Sbk2* and *Myl7*. 38 non-chamber selective genes were also upregulated in control samples and linked to enhancers by TBX5-dependent looping. These included *Gja1* and *Tead1*. Only 5 vCM-selective genes were found near control anchors. **c**, 50 genes neighbored KO anchors. These genes included 5 aCM-selective genes and 3 vCM-selective genes. Most of the differentially expressed genes near KO anchors were more highly expressed in Myocyte_6 (KO) compared to Myocyte_1 (Ctrl).
