## Supplementary material for "*Tbx5* maintains atrial identity by regulating an atrial enhancer network": Supp. Table 1. Summary of high throughput data

**Supp. Table 1. Summary of high throughputdata used in this study.**

| Multiome--snRNAseq | | | | | |
|---|---|---|---|---|---|
| Replicate | Nuclei | Sequencing Read Pairs; RNA | Valid barcodes (%) | Sequencing Read Pairs; ATAC | Valid barcodes (%) |
| Con1 | 4,207 | 414,649,927 | 96.6 | 213,071,387 | 97.2 |
| Con2 | 5,167 | 349,405,410 | 96.8 | 173,447,944 | 97 |
| KO1 | 2,254 | 364,314,050 | 95.3 | 196,443,737 | 97.2 |
| KO2 | 2,867 | 341,831,044 | 96.5 | 223,305,458 | 97.5 |

| HiChIP | | | | | |
|---|---|---|---|---|---|
| | Total PETS | % Reads in Anchors | % HQ Unique Mapped | % Long Range Interaction | % in Loops |
| Con1 | 436,618,010 | 38.74 | 80.97 | 3.3 | 0.25 |
| Con2 | 567,537,934 | 36.74 | 79.78 | 2.84 | 0.21 |
| KO1 | 589,331,106 | 27.71 | 82.19 | 4.23 | 0.19 |
| KO2 | 451,933,257 | 30.82 | 82.02 | 5.11 | 0.27 |

| TBX5 OE RNA-seq | | | | |
|---|---|---|---|---|
| | Total Properly Mapped | uniquely mapped | multi map % | % unmapped too short |
| con1 | 34,288,840 | 63.08% | 22.55% | 14.16% |
| con2 | 34,422,144 | 64.28% | 25.19% | 10.34% |
| con3 | 37,943,014 | 66.64% | 20.91% | 12.22% |
| con4 | 31,159,080 | 66.24% | 22.54% | 11.00% |
| con5 | 36,183,010 | 65.33% | 22.47% | 12.00% |
| TBX5OE1 | 32,907,046 | 67.70% | 17.71% | 14.35% |
| TBX5OE2 | 32,021,162 | 69.21% | 19.72% | 10.76% |
| TBX5OE3 | 33,677,240 | 67.78% | 18.79% | 13.14% |

| Other datasets | | |
|---|---|---|
| Description | GEO accession | Reference |
| Tbx5,TF, P300 aCM bioChip-seq | GSE215065 | Cao et al., Circulation, 2023 |
| aCM vs vCM RNAseq | GSE215065 | Cao et al., Circulation, 2023 |
