## Supplementary material for "*Tbx5* maintains atrial identity by regulating an atrial enhancer network": Supp. Table 7. Antibodies

**Supp. Table 7. Antibodies used in this study.**

| Antibody | Source | Catalogue No. | Host species | Western Dilution concentration | Immunofluorescence dilution |
|---|---|---|---|---|---|
| SAA | Sigma-Aldrich | A7811 | Mouse | - | 1:100 |
| FSD2 | Santa Cruz Biot. | sc-393072 | Mouse | | 1:100 |
| MYL7 | Proteintech | 60229-1-Ig | Mouse | | 1:50 |
| MYL2 | ab79935 | Abcam | Rabbit | | 1:100 |
| TBX5 | Abcam | 137833 | Rabbit | 1:500 | |
| GADPH | Invitrogen | PA1-16777 | Rabbit | 1:5000 | |
| mCherry | Abcam | ab167453 | Rabbit | | 1:100 |
